# PepCL: A replay-based continual learning framework for updating peptide-MHC models

**DOI:** 10.64898/2026.07.20.733558

**Authors:** Prathamesh M. Chati, Vishal D. Lashkari, Ankit Salhotra, Peter M. Bruno, Vasilis Ntranos

## Abstract

Understanding peptide-major histocompatibility complex (MHC) class I binding is critical for effective vaccine and immunotherapy design but is a combinatorially complex challenge for which prediction models have become essential. MHC ligands are typically identified at scale via untargeted mass spectrometry (MS), and this has built a strong base for peptide-MHC model training. However, MS incompletely captures the vast peptide-MHC space due to technical, sampling, and biological biases. Although recently developed experimental assays have queried such blind spots yielding complementary information, existing peptide-MHC predictors have not yet incorporated these orthogonal data and are not designed to be updated as new data are generated. Here, we introduce PepCL (<u>Pep</u>tide-MHC <u>C</u>ontinual <u>L</u>earning), a continual learning framework for updating peptide-MHC predictors with new assay data while explicitly preserving prior MS knowledge. To enable PepCL, we also develop MHCPrime, a new state-of-the-art pan-allelic peptide-MHC prediction model, trained on publicly available MS data, that can be effectively updated under our framework. We demonstrate that PepCL allows MHCPrime to learn previously unseen, assay-specific information while preventing catastrophic forgetting that is typically observed with conventional fine-tuning. We evaluate PepCL and MHCPrime in a variety of biological contexts, including infectious disease and cancer, and show improved peptide-MHC prediction that transfers across alleles for broader applicability in clinical settings. Overall, our results establish PepCL as a flexible framework for extending the utility of peptide-MHC models by improving their predictive performance as immunopeptidomics assays continue to evolve and new data become available.

## Introduction

Major histocompatibility complex (MHC) class I presents endogenous peptides to CD8^+^ T cells, mediating immune recognition of self and foreign proteins^1,2,3^. With more than 40,000 resolved alleles, each with distinct peptide binding preferences, the space of possible peptide-MHC interactions is too large to characterize experimentally at scale^4^. Computational prediction of peptide-MHC binding has therefore become central to epitope discovery, vaccine design, and neoantigen prioritization^5,6^. Most high-performing class I predictors are trained primarily on mass spectrometry (MS) eluted ligand (EL) data, often supplemented with binding affinity measurements^7,8,9,10,11^. This has been highly effective because MS provides the largest available view of peptides presented in cellular contexts and captures antigen processing^12,13^. However, MS-derived ligandomes are incomplete observations of all presentable peptides and reflect platform-, protocol-, and sampling-dependent biases, including underrepresentation of sequence classes such as cysteine-containing or highly hydrophobic peptides^14,15,16,17^.

Emerging experimental technologies are expanding the data available for understanding peptide-MHC binding and presentation beyond MS. Although these assays are not biologically equivalent to MS, they interrogate related determinants of peptide presentation, including peptide-MHC binding, complex stability, and allele-specific sequence preference^14,18,19,20^. Furthermore, these assays can provide experimentally evaluated non-binders, continuous readouts, and library designs that sample sequence regimes underrepresented in MS, specifically peptides outside the human proteome. Integrated with MS-derived ligands, this information can support predictors with broader coverage of peptide-MHC sequence space while retaining an anchor to endogenous presentation. Yet most existing models are distributed as fixed, versioned predictors and are not designed to be updated as new assay data become available. Incorporating new evidence typically requires full retraining, reimplementation of the original pipeline, or conventional fine-tuning (FT), none of which explicitly controls the trade-off between acquiring new assay-specific signals and preserving previously learned presentation knowledge^21,22^.

This trade-off is central because new peptide-MHC assays are not interchangeable measurements of a single biological endpoint^11^. MS eluted ligands, binding affinity measurements, display-based assays, and targeted presentation assays each sample related but distinct aspects of the binding-presentation process, with slightly varying binder/non-binder definitions, readout scales, and assay-specific biases. Direct fine-tuning of a model on these data can therefore improve performance on a target assay by shifting the model toward that assay’s experimental endpoint or selected peptide distribution, potentially overwriting MS-derived presentation information that remains valuable for applications within the cellular context (i.e. catastrophic forgetting)^23,24,25^. Prior work has shown that task-specific FT or transfer learning of peptide-MHC models can improve domain-specific predictions, supporting the broader value of updating models with additional experimental evidence; however, these approaches generally optimize only on the new target task without explicitly retaining pretrained MS knowledge^26,27^. Conversely, preserving the MS prior too strongly can prevent useful adaptation to new experimental signals. Thus, model updating requires a controlled framework that can absorb complementary information while limiting overspecialization to the incoming data distribution^28,29^.

Here, we introduce PepCL (<u>Pep</u>tide-MHC <u>C</u>ontinual <u>L</u>earning), a replay-based continual learning framework for updating peptide-MHC predictors with new assay data while explicitly controlling retention of prior MS domain knowledge. PepCL combines new data updating with replay from the original MS training distribution and drift regularization against a frozen pretrained model, allowing users to tune the balance between assay adaptation and retention of existing binding representations (**Fig. 1**)^30,31,32^. To provide an updatable backbone for our framework, we developed MHCPrime, an open-source, GPU-trainable pan-allelic transformer model trained on MS-derived ligands, designed for efficient customization, retraining, and repeated updating. We apply PepCL to MHCPrime and systematically evaluate updates across multiple assay modalities that have not previously been incorporated into peptide-MHC model training. Across a diverse range of peptide-MHC-related tasks, we show that replay-based continual learning enables robust incorporation of new peptide-MHC information while reducing the overfitting and catastrophic forgetting observed under conventional FT.

**Fig. 1:**
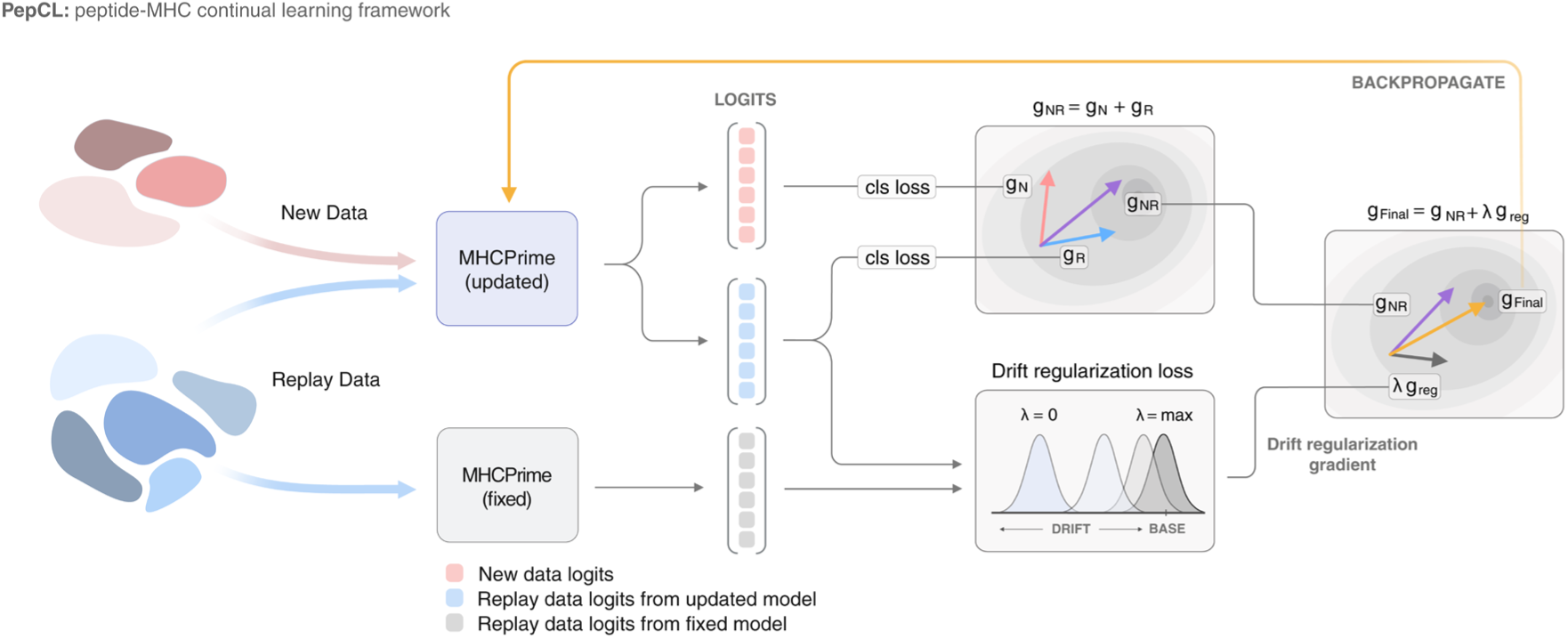
Overview of the PepCL framework. PepCL updates our pretrained MHCPrime model using both new data and replay data from MS. Two copies of the base model are first initialized: one to be updated and the other as a fixed prior. New and replay data batches are passed through the updated model, producing logits used to compute two separate classification losses (cls loss). The two gradients are then computed for the new (g_N_) and replay (g_R_) losses and summed to produce g_NR_. In parallel, the replay data are also passed through the fixed copy of the model. The replay data logits from the updated and fixed models are used to compute the drift regularization loss, which penalizes changes from the fixed model. The strength of drift regularization is controlled by a lambda parameter (λ), where greater values restrict the updated model more. We then compute the drift regularization gradient (g_reg_). Finally, the drift regularization gradient, g_reg_, and the summed classification gradient, g_NR_, are summed again to derive the final gradient, g_Final_, which is backpropagated through the updated model.

## Results

### MS-trained models underperform on non-MS data

We started by curating a broad set of peptide-MHC data spanning MS EL datasets along with four major non-MS assays: yeast display (YD), EpiScan, binding affinity (BA), and ESCAPE-seq (**Fig. 2a, Supplementary Fig. 1**)^8,14,19,20^. Along with capturing peptides from domains underrepresented in MS, including pathogen and cancer mutations, these assays provide experimentally defined weak- or non-binders, offering direct contrastive supervision that is not available in MS datasets. BA and ESCAPE-seq additionally provide continuous readouts that can be used to calibrate relative scores.

**Fig. 2:**
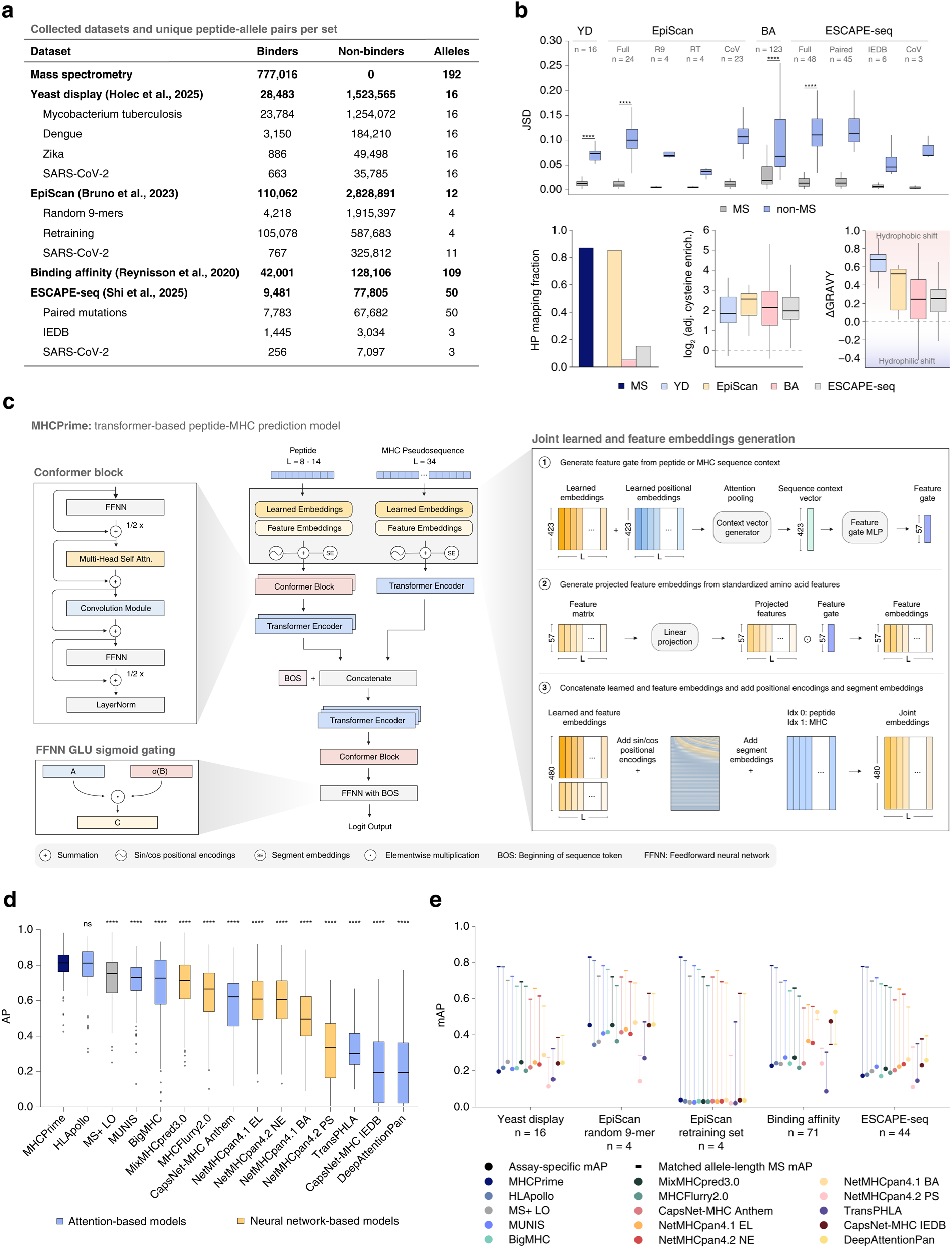
Dataset characterization and model benchmarking. **a)** Datasets collected and used for main updating experiments throughout this study. Binder and non-binder counts are unique at the peptide-allele level and represent filtered values. MS data paired against synthetic human proteome-derived negatives were used for pretraining the base model. Peptide-allele overlaps between MS and non-MS data were removed from non-MS sets before benchmarking (See **Methods** and **Supplementary Fig. 1)**. **b)** Sequence and biochemical differences between non-MS datasets and matched MS ligands. **Top**, position weight matrix (PWM) divergence between each dataset and MS, computed within matched allele- and peptide-length groups. For the reference (grey), 5% of peptides from each MS allele-length group were randomly sampled ten times; for each sample, a PWM was computed from the sampled peptides and compared by Jensen-Shannon divergence (JSD) to a PWM computed from the remaining MS peptides, then averaged across the ten repeats. For each non-MS dataset (blue), one PWM was computed from that dataset’s peptides within each matched allele-length group and compared by JSD to the corresponding MS PWM. Values above each dataset indicate the number of matched allele-length groups. P-values compare the non-MS JSD values with the matched MS reference values using paired Wilcoxon signed-rank tests. **Bottom left**, fraction of unique binder peptides mapping to the human proteome. **Bottom middle**, per-allele cysteine enrichment relative to MS. Values represent the log_2_ fold change of the cysteine enrichment ratio among non-MS assay binders versus their non-binder/background peptides relative to the corresponding enrichment ratio among MS ligands versus MS background peptides. **Bottom right**, per-allele ΔGRAVY, computed as the mean GRAVY score of non-MS binders minus the mean GRAVY score of matched MS ligands using the Kyte-Doolittle scale. **c)** MHCPrime model architecture. MHCPrime uses a dual-encoder architecture that separately encodes the peptide and MHC pseudosequence, then concatenates the representations while adding a beginning of sequence (BOS) token. Once moving through a series of joint transformer and conformer blocks, the BOS token is fed through the feedforward neural network (FFNN), which produces the final score. Diagrams for the conformer block, sigmoid gated linear unit (GLU) activation, and joint feature embeddings are in the outlined boxes. Additional model specifications can be found in the **Methods**. **d)** Benchmarking on the held-out MS test set. Average precision (AP) was computed separately for each of the 141 alleles using a 1:99 positive to negative ratio; boxplots show the distribution of per-allele AP values across models. Colors denote attention-based and neural network-based models. P-values were computed relative to the allele AP distribution of MHCPrime using paired Wilcoxon signed-rank tests. **e)** Benchmarking on non-MS datasets. For each assay set, points show assay-specific mean AP (mAP) across the available allele-length groups evaluated at a 1:99 positive to negative ratio. Connected rectangles show the corresponding matched MS reference mAP, computed on MS test samples restricted to the same allele-length groups as each non-MS dataset. The number of alleles in each dataset is shown on the x-axis. Significance labels indicate: ns, P > 0.05; *, P ≤ 0.05; **, P ≤ 0.01; ***, P ≤ 0.001; ****, P ≤ 0.0001.

Despite their smaller size relative to MS, non-MS assay datasets provide non-redundant information. This is evident at the peptide sequence level when comparing MS and non-MS binders, where the non-MS binders show significant Jensen-Shannon divergence (JSD) from MS across matched alleles and peptide lengths (**Fig. 2b, top**). Some of these representational shifts can be decomposed through sequence properties linked to known limitations of MS-derived datasets. For example, relative to MS, the non-MS assays contain peptides outside the human proteome with enrichment for cysteine-containing and more hydrophobic peptides (**Fig. 2b, bottom**). Though these analyses do not exhaust all assay differences, they show that the non-MS datasets occupy distinct regions of sequence space relative to MS and can potentially improve the representational diversity of MS-only training datasets.

We next built MHCPrime as the updatable pan-allelic backbone for PepCL. MHCPrime is a GPU-trainable, attention-based, peptide-MHC model designed to expose the components needed for continual learning, including direct access to model parameters, replay batches, custom losses, and repeated assay-specific optimization^33^. We designed MHCPrime following the architecture of HLApollo and CapsNet-MHC with several modifications to incorporate biochemical information and better capture local peptide context (**Methods**; **Fig. 2c**)^10,34^. To establish a general MS-trained reference point, we trained a single (non-ensembled) MHCPrime model on MS-derived ligands paired with synthetic negatives from the human proteome using a ranking-based classification loss. This yielded the base checkpoint used throughout this study. As a simple first-order baseline, we also evaluated an allele- and length-matched log odds model derived directly from the MS training binders^35,36^.

We benchmarked MHCPrime against held-out MS and non-MS datasets after removing peptide-allele overlaps to prevent train-test leakage. Following standard evaluation settings that reflect rare positive candidate screening, our classification benchmarks were evaluated at a 1:99 positive to negative ratio using average precision (AP), which is well-suited for highly imbalanced ranking tasks^10,16,37,38^. For BA and ESCAPE-seq, which contain continuous readouts, we also reported Spearman correlation evaluated with balanced binder/non-binder sets. Under these settings, MHCPrime achieves state-of-the-art performance on the MS benchmark, providing a strong baseline MS model (**Fig. 2d**). We observe a similar trend when evaluating on additional performance metrics (AUC, AUC0.1, and PPVn) per allele (**Methods**; **Supplementary Fig. 2**). Our results indicate that MHCPrime is able to perform on par with HLApollo, despite being a single, non-ensembled model trained only on mono-allelic data (**Methods**; **Supplementary Fig. 3**). Moreover, HLApollo is closed-source and only available as 10 ensembled models that must be run consecutively to yield predictions, making large-scale inference impractical. In contrast, MHCPrime is fully customizable, adaptable to new data, can accept any pseudosequence within the given length constraint, and is GPU-optimized. When evaluating the non-MS benchmarks, however, the performance of both MHCPrime and HLApollo declines substantially, a pattern also observed across other models trained primarily on MS data (**Fig. 2e**; **Supplementary Fig. 4**). Notably, methods trained with both MS and BA, such as NetMHCpan4.1, show better performance than MS-only models on non-MS data, supporting the idea that complementary assay signal can improve cross-domain behavior (**Supplementary Fig. 5**)^8^.

### Replay-based continual learning supports efficient model updating under representationally diverse MS data

Conventional fine-tuning (FT) updates a pretrained model using only the incoming dataset, so every gradient step is driven by the new data distribution. PepCL instead updates the model while preserving access to the MS prior through two mechanisms: data replay and base model drift regularization^30,31,32^. Replay reintroduces previously observed MS peptides during updating, providing gradients that counterbalance the new data distribution and reduce loss of prior domain performance. Drift regularization adds a second constraint by comparing the updated model with a frozen copy of the pretrained model on replay peptides and penalizing changes in their raw scores using mean squared error (MSE)^39^. The strength of drift regularization is controlled by a parameter, lambda (λ), where λ = 0 corresponds to replay-only updating and greater values increasingly anchor the updated model to the pretrained model. The MS replay and new data classification losses, computed with the same ranking-based classification objective used to train MHCPrime, are combined with the drift regularization loss for the final model update step. To characterize these components before applying PepCL to non-MS assay datasets, we performed a series of controlled experiments with MS data.

We first considered in-distribution updates, where base models were first pretrained on increasing fractions of MS data and then updated with an additional 10% of held-out MS data. Since early checkpoints trained on small MS fractions are intentionally undertrained, we used PepCL with replay-only (λ = 0) in this experiment rather than constraining the update to preserve undertrained predictions. To determine whether updating could approximate full retraining, we performed updates with 1,000 steps. As expected, increasing the amount of MS pretraining data improved performance but showed diminishing gains at higher pretraining data fractions, suggesting that simply adding more in-distribution MS data eventually provides limited additional benefit without greater representational diversity (**Fig. 3a**). In this relatively ideal setting, where the new data were drawn from the same MS distribution, PepCL recovered most of the gain obtained by retraining while avoiding the performance loss seen with FT (**Fig. 3b**). Although replay introduces additional forward passes relative to FT, both approaches receive the same number of gradient updates; even after accounting for this overhead, PepCL remained substantially less computationally expensive than full model retraining (**Fig. 3c**).

**Fig. 3:**
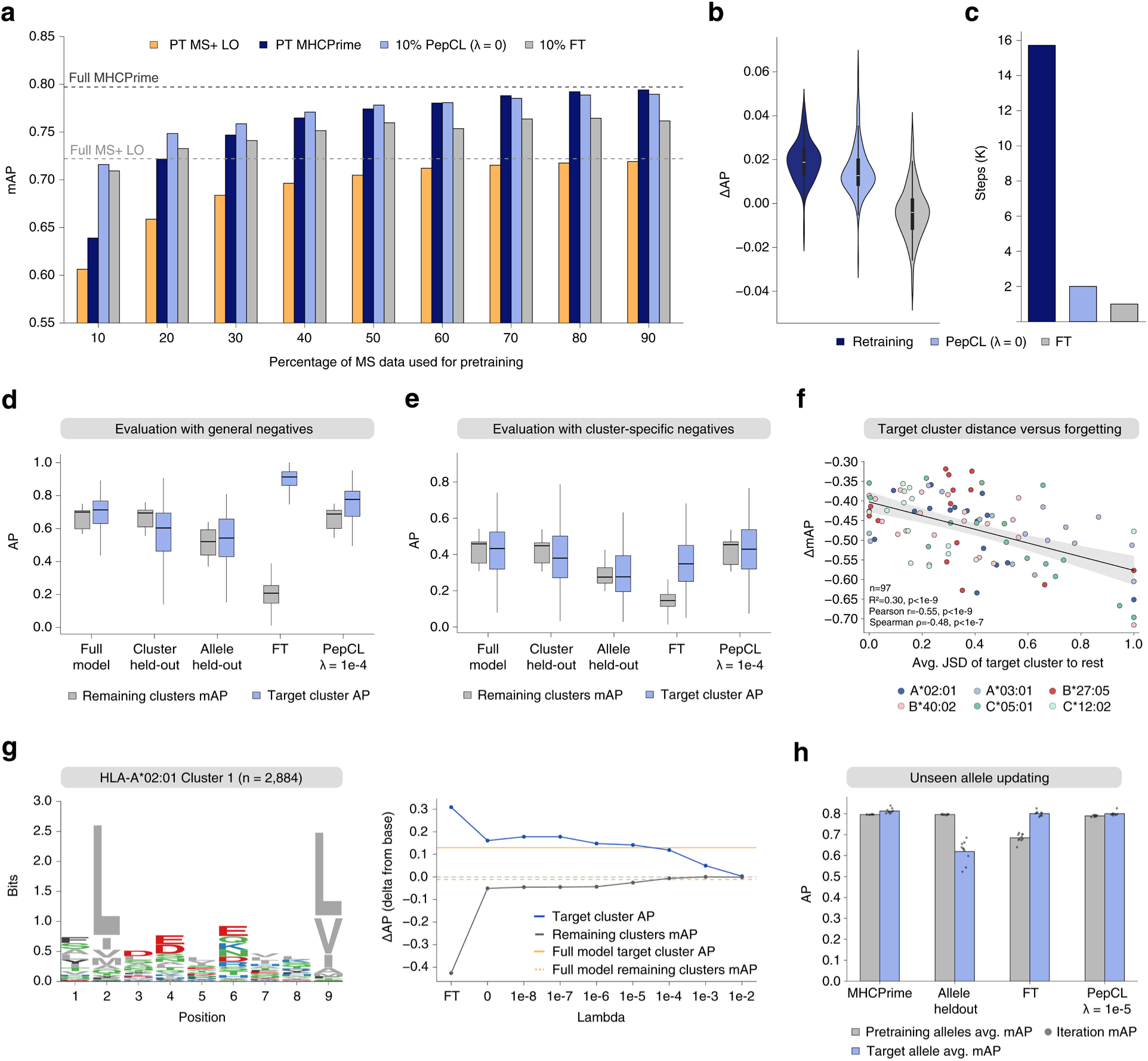
Controlled MS updates define adaptation and retention trade-offs. **a)** MS mAP across 142 alleles after pretraining on increasing fractions of MS data and updating with an additional 10% MS subset. Yellow bars show log odds performance for the current subset of data (PT: pretrained; MS+ LO: MS binder log odds), and navy bars show performance after pretraining on the same subset; light blue and grey bars show performance after updating the corresponding pretrained model with 10% additional data using either PepCL (λ = 0) or FT for 1000 steps, respectively. Dashed lines indicate the full MS model (MHCPrime) and full MS log odds derived from all MS training binders. **b)** Per-allele AP difference (ΔAP) after adding the next 10% MS subset. For each allele, ΔAP is computed relative to the corresponding pretrained checkpoint, then averaged across the 9 subsets. **c)** Training steps required for full retraining, PepCL, and FT. Steps measure the number of forward passes. **d)** Target cluster holdout benchmark evaluated with general human proteome negatives. Light blue distribution shows AP per target cluster; grey distribution shows mAP of all remaining clusters excluding the target cluster. The full model is the pan-allelic model retrained with all alleles and lengths, with the target allele containing 9-mers only; the cluster held-out shows models where each cluster is held out from training; the allele held-out shows the full model excluding the target allele from training. **e)** Same evaluation setting as in d), but cluster-specific binders are evaluated against cluster-specific negatives. **f)** Relationship between the remaining cluster performance difference after FT and the target cluster distance. For each allele-cluster holdout, the x-axis shows the average JSD between the target cluster and all remaining clusters, computed from positive binders; the y-axis shows the mAP of remaining clusters for the full model minus the mAP of remaining clusters after FT, excluding the target cluster. The black line shows a fitted linear regression, and the grey band indicates the 95% confidence interval. Reported values indicate R^2^, Pearson correlation, and Spearman correlation (n = 97). **g) Left**, sequence logo plot for the representative holdout cluster from HLA-A*02:01 using 2,884 cluster-specific binders. **Right**, the target cluster AP difference from the full model and the mAP difference of remaining clusters from the full model for FT and across various lambda values. **h)** Unseen allele updating benchmark using fixed downstream PepCL settings (λ = 1e-5; 200 steps).

We next tested a more challenging setting in which the update distribution was intentionally shifted from the pretraining distribution. For six major alleles, we performed clustering on feature embeddings of MS 9-mer ligands to derive motif-defined groups, trained base models with each cluster held out, and then updated the models with the held-out cluster while using allele-specific peptides from the remaining clusters as the replay set. This created a controlled test of whether updating could recover a previously unseen motif regime without degrading motifs already learned for the same allele. For these updates, we used a shorter fixed 200-step budget and introduced drift regularization with λ = 1e-4. We then used AP to measure the performance on the cluster being updated and the mean AP (mAP) of all remaining clusters to evaluate the performance of the prior. Conventional FT produced the strongest recovery on the target cluster when evaluated against a general human proteome-derived negative background, but this came at a severe cost to the remaining clusters, in some cases reducing performance below that of the model in which the target cluster had been held out (**Fig. 3d**; **Supplementary Fig. 6**). This represents a controlled example of catastrophic forgetting: the model rapidly adapts to the new motif but loses broader allele-specific knowledge. By contrast, PepCL improved performance on the held-out cluster while preserving performance on the remaining clusters, indicating a more balanced update. We repeated our evaluation using cluster-specific human proteome-derived negatives, asking whether each update strategy had learned a nuanced boundary within the missing motif region rather than simply exploiting broad motif-level features. Under this harder evaluation, PepCL achieves the strongest cluster-specific performance and closely matches the full reference models. Our results suggest that replay-based updating better preserves within-motif separability, while FT learns a less discriminative boundary by overspecializing to the cluster-specific motifs (**Fig. 3e**).

To further explore determinants of catastrophic forgetting, we returned to the original evaluation setting and found that forgetting under FT is distance-dependent: updates from motifs farther from the clusters used during pretraining produce larger losses across those remaining clusters (**Fig. 3f**). This aligns closely with our prior findings, where in-distribution updates under FT caused substantially less degradation than we observe in our cluster-specific updates. Lastly, to demonstrate how the drift regularization lambda (λ) parameter can enable finer control over cluster- and prior-specific performance, we performed a sweep over λ values for a representative motif cluster from HLA-A*02:01 (**Fig. 3g**). Replay provides the first major reduction in catastrophic forgetting of the prior and overfitting of the cluster, with higher λ values then allowing for more granular adjustments to reach the full model performance. Because the cluster holdout test represents an intentionally severe setting, we originally used a λ value of 1e-4. However, in most settings, incoming data are expected to contain a mixture of familiar and novel peptide motifs; we therefore used a single moderate setting, λ = 1e-5, and a fixed 200-step budget for all remaining analyses in this study.

Finally, we extended this analysis to unseen alleles, where updating with our default settings introduced allele-specific information absent from the base model. In this setting, PepCL again improved held-out allele performance while maintaining performance on the remaining alleles, showing that replay-based updating can incorporate new allelic contexts without the stronger degradation observed under FT (**Fig. 3h**).

These controlled experiments establish PepCL as a balanced updating strategy that allows for computationally efficient incorporation of new information into our pretrained peptide-MHC model while limiting overspecialization and catastrophic forgetting using replay and drift regularization. The benefit becomes most apparent when the incoming data are distributionally distinct from the original training set.

### PepCL enables learning assay-specific signals while retaining pretrained MS domain knowledge

We next applied PepCL to update our base model with several assay-specific datasets, beginning with the Yeast Display (YD) data^19^. Consistent with our synthetic experiments, both FT and PepCL significantly outperform the base model on the YD benchmark set, but FT suffers a disproportionately larger loss on the MS benchmark (**Fig. 4a**). This trade-off becomes clearer at the per-allele level, where PepCL generally retains most of the YD gain while preserving substantially more MS performance than FT (**Fig. 4b**). Similarly, when benchmark sets were constructed as mixtures of MS and YD data, FT performs best only near the pure YD end, while PepCL achieves stronger performance across a broader range of assay compositions (**Fig. 4c**). We further evaluated ranking and recovery on a set of MS-derived peptides across the pathogen domains in the YD sets and observed that PepCL either maintains or improves current recoverability at the traditional 2% threshold while FT often diminishes performance on this benchmark (**Supplementary Fig. 7**). Lastly, we performed updates in the same manner with ESCAPE-seq cancer mutation data and evaluated on cancer data from Ligand.MHC Atlas (**Supplementary Fig. 8**). We observed both PepCL and FT improve the recovery and ranking of reported tumor neoantigens, however, only PepCL was able to improve the rankability of peptides MS-derived from Ligand.MHC Atlas. This demonstrates that replay-based continual learning provides a more balanced adaptation to the new assay than conventional FT alone and that specialization observed in FT does not carry to related domains in MS data.

**Fig. 4:**
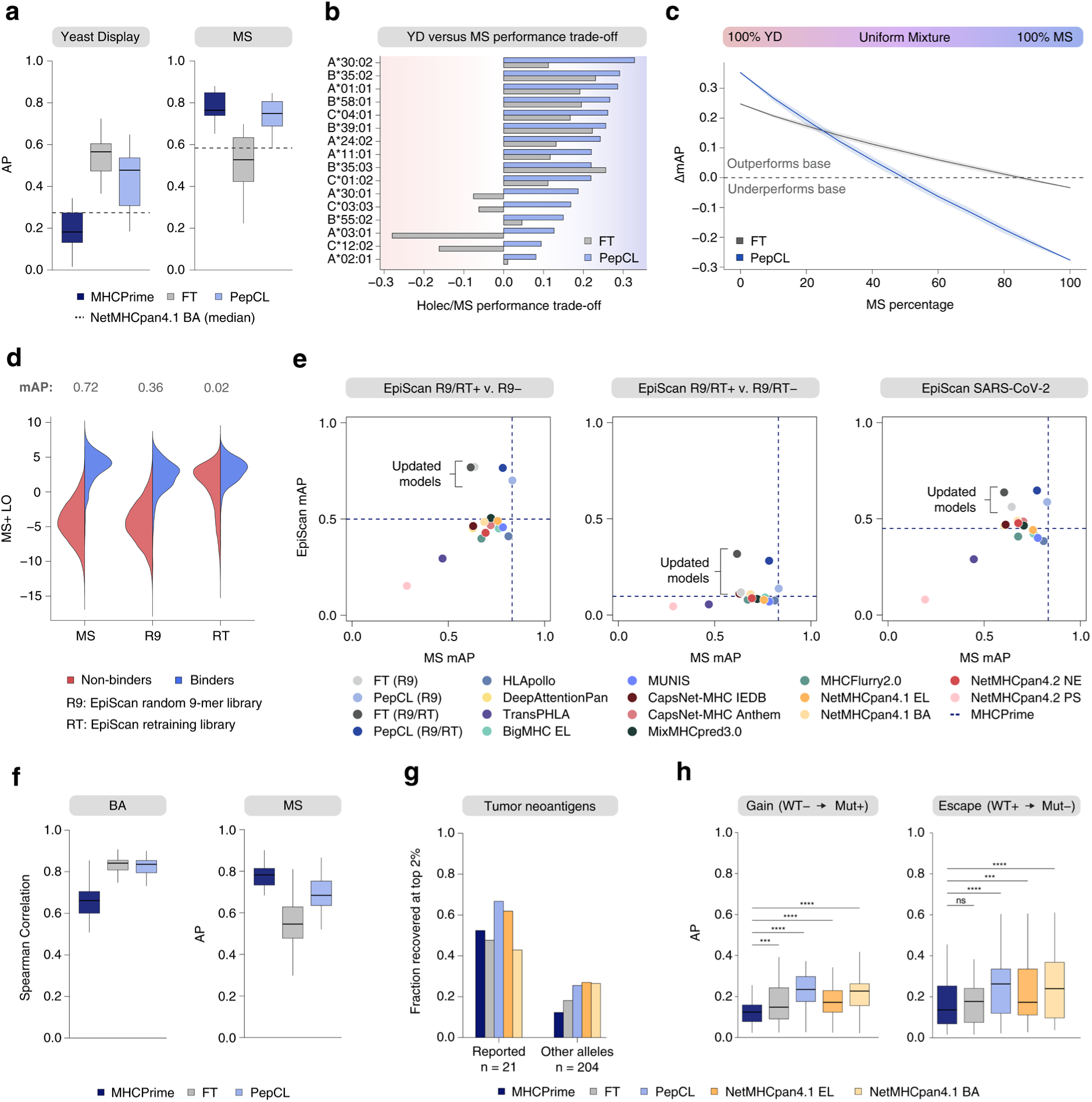
PepCL can integrate assay-specific readouts across different tasks. **a)** YD and MS test set AP held at a 1:99 positive to negative ratio after updating on YD data across 16 alleles. The MS test set was matched to allele-length groups within the YD data. Dashed lines indicate NetMHCpan4.1 BA. **b)** Per-allele YD/MS performance trade-off after updating. Net benefit is computed as YD gain relative to MHCPrime minus any MS loss relative to MHCPrime; positive values indicate YD improvement after accounting for MS degradation. **c)** Performance across mixed YD/MS evaluation sets. Each point represents mAP change relative to MHCPrime for evaluation mixtures spanning 100% YD to 100% MS in increments of 10%. **d)** MS binder log odds scores for MS ligands and EpiScan R9 and RT binders/non-binders. mAP values above each distribution summarize binder/non-binder separability. **e)** EpiScan benchmark performance after updating with R9− or R9/RT− negative backgrounds. Points show EpiScan mAP and matched MS mAP for public models, MHCPrime, FT, and PepCL across the EpiScan R9/RT+ versus R9−, R9/RT+ versus R9/RT−, and SARS-CoV-2 benchmarks. Dashed lines indicate MHCPrime performance. The EpiScan mAP values were generated by sampling the corresponding datasets at a 1:99 positive to negative ratio per allele 100 times, then averaged. **f)** Continuous BA updating. **Left**, Spearman correlation on BA test set held at a 1:1 positive to negative ratio; **right**, AP on the matched MS test set held at a 1:99 positive to negative ratio after BA updating. **g)** Tumor neoantigen recovery within the top 2% of rank values using the models updated on the ESCAPE-seq continuous E-scores shown for reported neoantigen allele assignments and alternative allele pairings. **h)** Differential mutation scoring in gene-held-out ESCAPE-seq experiments. AP is shown for gain and escape mutations, using model-predicted mutant to wild-type score differences. P-values are computed relative to the MHCPrime gene AP distribution using paired Wilcoxon signed-rank tests. Significance labels indicate: ns, P > 0.05; *, P ≤ 0.05; **, P ≤ 0.01; ***, P ≤ 0.001; ****, P ≤ 0.0001.

We continued our evaluation of PepCL by updating with EpiScan data, which contains two primary libraries: a random 9-mer library (R9), which provides a broad background of mostly easy negatives (peptides that lack canonical motifs), and a retraining library (RT), which was intentionally enriched for peptides predicted to bind by existing models and therefore yields a substantially harder negative background^14^. This distinction is evident from the MS binder log odds (**Fig. 4d**). Whereas binders and non-binders in the R9 library remain partially separable, the RT binders and non-binders exhibit extensive overlap in MS space and near-random ranking performance, indicating that many RT negatives retain canonical binding features and cannot be rejected by simple anchor-based heuristics alone that most existing models rely on. Therefore, EpiScan data provide a unique opportunity to train on hard negatives, which is difficult to achieve with synthetic negatives because they may include unobserved true binders. We constructed two update sets with the same mixed binder pool (R9/RT+) but different negative backgrounds, using either random negatives (R9−) or a combined hard negative background sampled from both libraries (R9/RT−). On the easier R9− benchmark, both update methods perform well (**Fig. 4e, left**). In contrast, improvement on the harder R9/RT− benchmark requires exposure to hard negatives themselves: models trained with only R9− show little benefit, whereas those trained with R9/RT− achieve clear gains (**Fig. 4e, middle**). The same pattern extended to the SARS-CoV-2 EpiScan benchmark, where models trained with the hard R9/RT− background perform best (**Fig. 4e, right**). Thus, experimentally validated hard negatives provide discriminative information that is not recovered from easy random negatives alone. Furthermore, replay-based continual learning often matches or exceeds FT in absorbing this signal.

Finally, we tested whether PepCL could help integrate continuous value readouts. We first updated our base model with BA measurements using a continuous ranking objective while maintaining the original classification loss on the replay set and observed improved relative ordering of peptides by binding strength relative to the base MHCPrime (**Fig. 4f**; **Supplementary Fig. 9**)^8^. We next used the continuous E-score values from ESCAPE-seq, a measure of peptide-MHC cell surface display, across two tasks: traditional threshold-based recovery of reported tumor neoantigens and threshold-free differential mutation scoring^20^. In the first recoverability task, FT and PepCL improve recovery of reported tumor neoantigens, with PepCL performing best overall (**Fig. 4g**). We then performed a gene holdout experiment, systematically excluding each target gene’s peptides during updating and evaluating it using model-predicted score differences between paired mutant and wild-type peptides (Δ = **S_Mut_** - **S_WT_**). In this framework, gain mutations (WT−/Mut+) are expected to receive strongly positive values, whereas escape mutations (WT+/Mut−) are expected to receive strongly negative values, providing a threshold-independent approach to ranking mutation-driven presentation changes. Under this setting, PepCL shows a much larger advantage than FT in prioritizing both gain and escape mutations, indicating that replay-based continual learning better preserves the relative score structure needed to traverse nearby sequence variants and identify mutations that most strongly perturb presentation (**Fig. 4h**; **Supplementary Fig. 10**).

Overall, our results establish the utility of PepCL in expanding the predictive power of MS-trained peptide-MHC models across a diverse pool of assays and tasks, with replay-based continual learning offering the most robust path to incorporating new signals without assay-specific drift.

### Pan-allelic continual learning transfers assay-derived signals to unseen alleles

A practical challenge for incorporating emerging peptide-MHC assays is that most profile only a small subset of common alleles. Consequently, an effective updating framework should not only improve the alleles directly observed during training but also transfer newly acquired information across the broader pan-allelic sequence space^40,41^. Unlike the assay adaptation experiments above, this setting requires the model to propagate information through shared peptide-MHC representations rather than simply prioritizing alleles present in the update set. We therefore asked whether assay-derived signals learned during updating could improve prediction for alleles not included in the update set itself.

We first evaluated this using EpiScan SARS-CoV-2 data, where several alleles were absent from the update set but remained available for independent evaluation. For both the R9− and R9/RT− trained models, updating on only the four profiled alleles with PepCL consistently improved prediction on these previously unseen alleles, reducing the number of cases where allele-level performance fell below the base MHCPrime model (**Fig. 5a**). Furthermore, this demonstrates that pan-allelic representations learned by MHCPrime can enable assay-specific information to propagate beyond directly observed peptide-allele pairs.

**Fig. 5:**
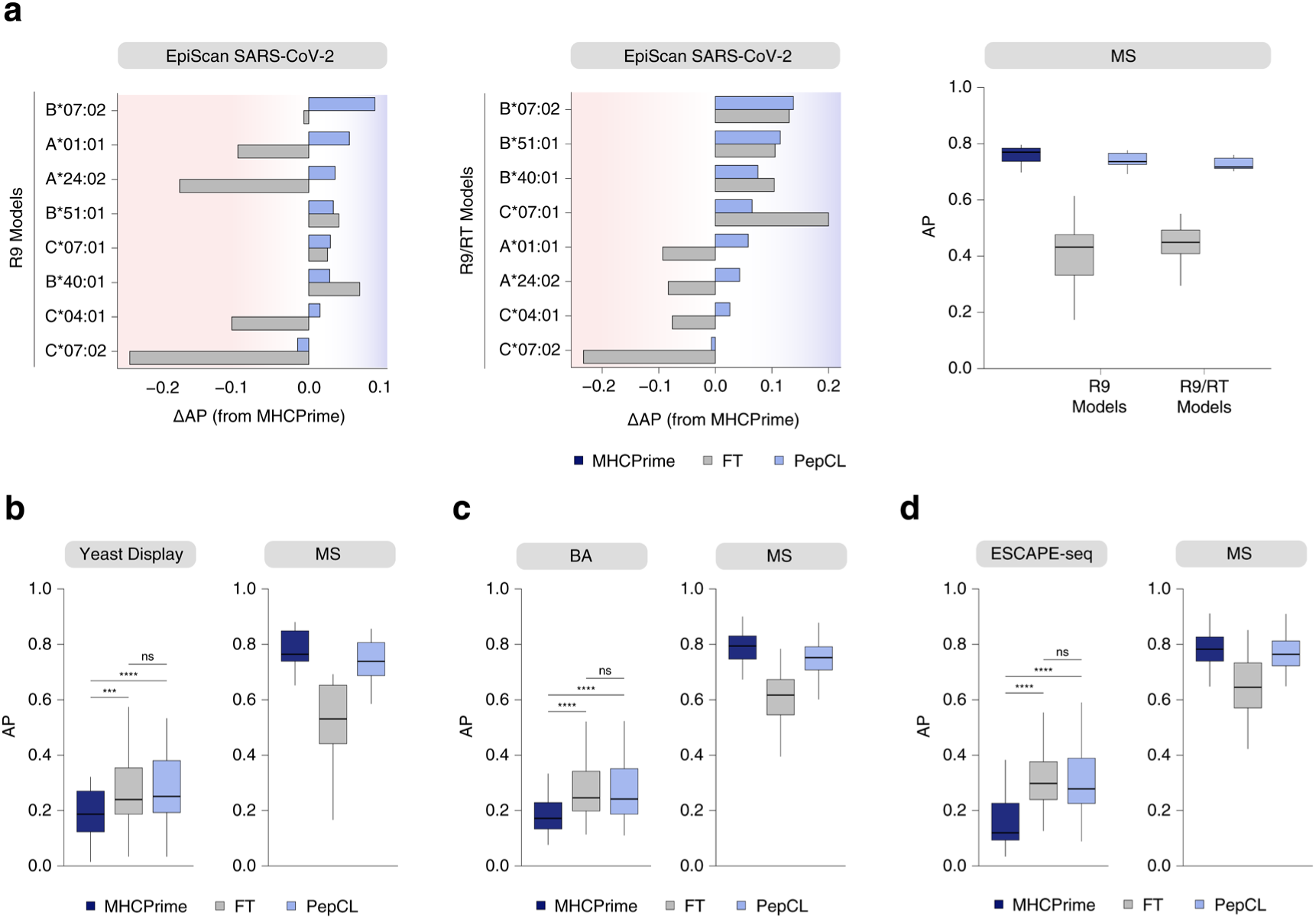
Cross-allele transfer after assay-specific updating. **a)** EpiScan SARS-CoV-2 transfer to alleles not observed during updating. **Left** and **middle**, per-allele AP differences relative to MHCPrime for unseen alleles after updating on the observed EpiScan alleles using R9− or R9/RT− training sets. **Right**, matched MS test set AP for the same held-out alleles after updating. **b)** Systematic allele holdout transfer on YD data. For each target allele, the allele was removed from the update and replay sets, and models were updated using the remaining alleles**. Left**, held-out YD AP; **right**, matched MS AP for the same alleles. **c)** Same test and evaluation as b) for BA data. **d)** Same test and evaluation as b) for ESCAPE-seq data. P-values are computed across allele AP distributions using paired Wilcoxon signed-rank tests. Significance labels indicate: ns, P > 0.05; *, P ≤ 0.05; **, P ≤ 0.01; ***, P ≤ 0.001; ****, P ≤ 0.0001.

To determine whether this behavior generalized beyond a single dataset, we performed systematic allele holdout experiments across the YD, BA, and ESCAPE-seq datasets. For each target allele, the allele was excluded entirely from both the assay-specific update set and replay set, and the model was updated using the remaining alleles with the same classification loss applied to both the new and replay datasets before evaluating on the held-out allele (**Methods**). Across the three datasets, FT and PepCL improved held-out allele performance over the base model, indicating that assay-derived signals can transfer across alleles even when the target allele is not directly observed during updating (**Fig. 5b-d**). However, the two update strategies differed in their effect on the original MS domain. FT produced comparable transfer gains but caused larger losses on MS benchmarks for the same alleles. In contrast, PepCL maintained more of the pretrained MS knowledge while achieving similar held-out assay performance. Thus, replay-based continual learning can extend assay-derived signals to unseen alleles without the same degree of loss in the prior domain.

### PepCL conservatively adopts assay-specific biases by anchoring to MS representations

Since FT and PepCL differ in how strongly they remain anchored to prior MS-derived binding preferences, comparing their learned motif changes provides a practical way to distinguish stronger assay-specific specialization from more constrained adaptation. Here, we illustrate this capability using assay-specific features documented in the underlying experimental systems.

We first examined EpiScan, where several peptide-level shifts relative to MS have been previously described^14^. Using models updated on the harder R9/RT− background, we scored 100,000 peptides from the human proteome and derived PWMs from the top-scoring peptides for each allele. Relative to the base model, both FT and PepCL increase recovery of cysteine-containing peptides and shift predictions toward greater hydrophobicity, consistent with sequence properties that are underrepresented in MS-derived ligandomes (**Fig. 6a**). However, FT generally showed a larger shift, while PepCL retained a more tempered departure from the MS prior. We next examined two additional documented EpiScan effects in HLA-A*02:01 and HLA-B*08:01, both of which highlight how assay-linked preferences can differ in how readily they should be absorbed.

**Fig. 6:**
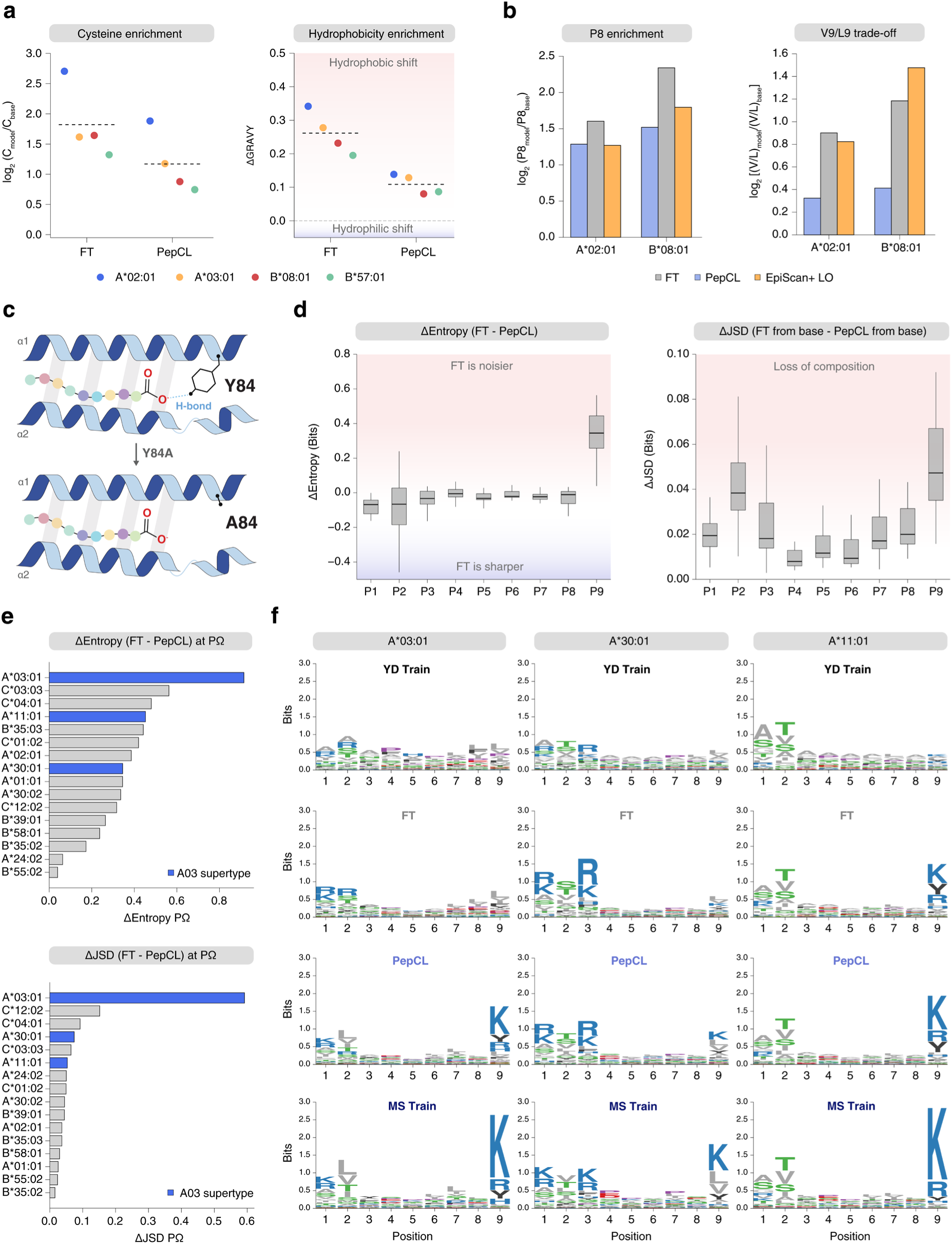
Sequence preference shifts differently between FT and replay-based updating. **a)** Cysteine and hydrophobicity shifts among top-scoring peptides after updating with EpiScan data. For each allele, 100,000 random human proteome-derived peptides were scored with MHCPrime, FT, and PepCL models updated on the EpiScan R9/RT− hard negative background. The top 2% of peptides for each model were used for analysis. **Left**, log_2_ fold change in cysteine-containing peptides relative to MHCPrime. **Right**, change in mean GRAVY score relative to MHCPrime, computed using the Kyte-Doolittle scale. Dashed lines indicate the mean across alleles. **b)** Penultimate proline enrichment and C-terminal valine/leucine (V/L) preference among top-scoring peptides. For each allele, the top 2% of scored human proteome-derived peptides were compared with MHCPrime. **Left**, log_2_ fold change in peptides containing proline at the penultimate position. **Right**, log_2_ fold change in the ratio of C-terminal V to L among top-scoring peptides ending in either V or L. EpiScan+ LO denotes the log odds derived from EpiScan R9/RT+ binders and applied to the same human proteome background. **c)** Schematic of the Y84A substitution in the YD single-chain trimer (SCT) system used by Holec et al., illustrating the altered C-terminal pocket near the peptide P9 position. **d)** Positional entropy and JSD differences between FT and PepCL after updating MHCPrime with YD data. For each allele, 100,000 random Swiss-Prot peptides were scored with the FT and PepCL models, and PWMs were generated from the top 2% highest scoring peptides. **Left**, positional entropy difference, computed as FT minus PepCL. **Right**, positional JSD difference from the base MHCPrime model, computed as JSD(FT, MHCPrime) minus JSD(PepCL, MHCPrime). Each point in a box represents one allele (n = 16). **e)** Allele-level P9 differences between FT and PepCL from the analysis in d). **Top**, P9 entropy difference, computed as FT minus PepCL. **Bottom**, P9 JSD difference from the base MHCPrime model, computed as JSD(FT, MHCPrime) minus JSD(PepCL, MHCPrime). Blue bars indicate alleles in the A03 supertype. **f)** Sequence logos for selected A03 supertype alleles. Logos show the YD training binders, the top 2% highest scoring peptides from FT and PepCL models, and the MS training binders for HLA-A*03:01, HLA-A*30:01, and HLA-A*11:01.

First, EpiScan showed increased penultimate proline enrichment, a more context-dependent shift because proline-adjacent peptides can be underrepresented in endogenous MS datasets by proteasomal processing rather than binding itself; both methods moved toward this signal, with PepCL again shifting more moderately than FT (**Fig. 6b, left**)^42^. By contrast, the valine versus leucine trade-off at P9 is more difficult to interpret as broadly desirable and likely reflects stronger platform-specific preference; here again, PepCL limited the shift relative to FT (**Fig. 6b, right**). Taken together, these shifts suggest that replay-based continual learning incorporates assay-derived signal through a stronger prior MS anchor, while conventional FT more readily absorbs the full distributional imprint of the new data, including both assay-linked preferences and dataset-specific sampling effects. This difference becomes especially important when the assay contains a documented perturbation that directly reshapes peptide preference rather than simply exposing underrepresented signals.

One example is the Y84A mutation made in the YD system from Holec et al., where prior work has shown that the single-chain trimer (SCT) design can relax native constraints at the peptide C terminus (**Fig. 6c**)^43^. In native class I complexes, Y84 supports the peptide C terminus through steric enclosure and hydrogen bond coordination of the P9 residue and terminal carboxyl group at the F pocket; alanine substitution at this position is therefore expected to weaken these interactions and broaden P9 permissiveness^44,45,46,47^. We therefore treated P9 as the most interpretable axis along which this perturbation should be visible. To examine how this feature was reflected in learned binding representations, we scored 100,000 peptides from Swiss-Prot with FT and PepCL and constructed PWMs from the top-scoring peptides for each allele. Across all alleles, P9 showed the clearest increase in entropy under FT relative to PepCL, while the corresponding increase in JSD was more modest, indicating that the dominant effect was not only replacement of residue preference but a broader, less restrictive terminal anchor under conventional FT (**Fig. 6d**). Although this does not establish Y84A as the sole driver of the observed shift, the concentration of the strongest signal at P9 is consistent with prior evidence that the SCT mutation relaxes C-terminal anchoring. This effect was most pronounced in HLA-A*03:01, which motivated closer inspection of related alleles, including HLA-A*30:01 and HLA-A*11:01 (**Fig. 6e**)^48^. From the sequence logo plots, the YD data showed weakened or redistributed terminal anchoring relative to the MS training distribution, and FT more fully absorbed this shift. In contrast, PepCL retained a stronger MS-like P9 preference while still partially adapting toward the assay-derived motif (**Fig. 6f**). Thus, replay-based continual learning did not block uptake of the YD signal, but tempered the most pronounced C-terminal deviations seen under unconstrained fine-tuning.

The EpiScan and YD examples show that not all departures from the MS prior carry the same interpretation. Some may reflect binding information that is underrepresented in MS, while others more plausibly reflect assay- or dataset-specific effects. By contrasting unconstrained and replay-based updating, we observe that PepCL absorbs new sequence preferences more conservatively and that they remain strongly shaped by the prior MS-derived binding structure.

## Discussion

Peptide-MHC predictors have become central tools for prioritizing candidate presented peptides across antigen discovery, vaccine design, and cancer neoantigen selection. However, many of the existing predictors are trained solely on MS data, limiting their applicability to domains beyond those captured by MS. The development of new assays that query related information to peptide-MHC presentation, such as YD, EpiScan, BA, and ESCAPE-seq, has added a large volume of distributionally diverse data to the training pool. These non-MS datasets contain valuable information that can naturally serve as an extension to MS datasets. Yet, many of these datasets are rarely integrated with current models and doing so requires retraining or fine-tuning. While retraining can serve as a reasonable first-pass, performing proper sampling and balancing of new and prior data is challenging, and can often lead to the larger dataset, MS, dominating the smaller. Furthermore, retraining can be cumbersome when the models are larger, which can hinder iterative testing. A faster option is to fine-tune the model with the new data; however, this approach can often lead to overspecialization on both dataset- and assay-specific properties while forgetting the prior training knowledge.

To address these issues, we introduced PepCL, a replay-based continual learning framework that combines GPU-efficient model updating with pretraining sample replay and drift regularization to anchor updates to the prior MS knowledge. PepCL was designed from principles established in continual learning and multi-task optimization, where model updates must balance plasticity to new data with stability on previously learned tasks^49^. Rehearsal and experience replay methods reduce forgetting by interleaving examples from previous tasks while learning the new task. Additionally, functional regularization and distillation-based methods constrain the updated model to preserve behavior from a previous frozen model state. PepCL combines these ideas for peptide-MHC prediction by replaying allele-specific samples from the MS training distribution while adapting to a new data set, with drift regularization against a frozen copy of the starting model.

Along with PepCL, we introduced MHCPrime, a GPU-efficient, transformer-based peptide-MHC model designed to serve as the updatable backbone for our framework. We developed MHCPrime as opposed to using an existing published model because PepCL requires direct access to model parameters, logits, losses, gradients, frozen model copies, and replay batches, which are necessary for testing and debugging each component of PepCL in a controlled way. The strongest transformer-based comparator in our benchmarks, HLApollo, provides a highly competitive reference point but is built as a closed-source, ensembled workflow. Our goal required a single, modular, fully retrainable model that could be repeatedly updated across new objectives. Architecturally, MHCPrime was designed to combine recent peptide-MHC modeling strategies with additional local peptide context and biochemical features. Despite being trained only on mono-allelic MS data and evaluated as a single non-ensembled model, MHCPrime outperformed all benchmarked models and competes directly with the fully ensembled HLApollo model. Thus, MHCPrime provides both a state-of-the-art MS-trained starting point and the practical model backbone needed for PepCL.

With PepCL implemented on MHCPrime, our results show that conventional FT and PepCL occupy different points along the adaptation-retention trade-off. FT often moved more strongly toward the incoming dataset, but this came with greater overspecialization and loss of prior MS performance. In the synthetic cluster-specific experiments, this was directly observable as FT strongly improved cluster-specific performance but degraded the pretrained knowledge of other clusters from the same allele. In assay-specific settings, the same phenomenon is harder to define because the incoming data differs by experimental endpoint, negative construction, readout scale, and sequence distribution. However, when updated models were evaluated on matched MS benchmarks and related MS-derived domains, FT often reduced performance, while PepCL generally preserved or improved the base MS model. This suggests that FT can learn useful assay signals while also absorbing dataset- or assay-specific biases that may not translate cleanly back to endogenous presentation. Analysis of the model features supports this interpretation. For both EpiScan and YD, conventional FT fully absorbed documented dataset- and assay-specific biases, while PepCL retained a more balanced profile but did not completely block adaptation to new information.

This distinction is central to how updated peptide-MHC models should be used. If the goal is to optimize predictions for a specific assay, conventional FT can be appropriate and may produce the strongest assay-specific model. However, many external assays are designed as proxies for processes that ultimately occur in cells, including binding, stability, surface display, and presentation. In that setting, the goal is not simply to reproduce the assay distribution, but to incorporate the information it provides while retaining MS-derived presentation knowledge. PepCL is designed for this use case. It provides a more conservative update, which is useful as preservation of the MS prior is important for most downstream applications, including the design and development of immunotherapies such as therapeutic cancer vaccines^50,51,52^.

Beyond assay-specific gains on the alleles directly observed during updating, the pan-allelic structure of MHCPrime allowed assay-derived information to transfer to alleles absent from the update set. This capability addresses a structural limitation of the assays themselves: throughput constraints mean that most peptide-MHC assays profile only a handful of common alleles, even though the alleles that matter clinically span a far larger and more diverse space^53^. That space is also unevenly characterized as publicly available ligand and assay data are concentrated on common, predominantly European-ancestry alleles, while many alleles prevalent in African, Asian, and other under-sampled populations remain sparsely measured^54^. A framework that propagates assay-derived signal through shared peptide-MHC representations therefore extends the practical value of a limited-allele assay across the broader allelic landscape, which is a prerequisite for epitope and vaccine prediction that generalizes across populations rather than only the alleles an assay happened to include^55^. In the allele holdout experiments, both FT and PepCL improved performance on unseen alleles, confirming that shared representations can carry assay-derived signal across alleles. Critically, PepCL achieved this transfer with substantially less degradation on matched MS benchmarks, indicating that replay-based continual learning is particularly well suited to extending narrow-panel assay data into broad pan-allelic prediction.

Lastly, there are still limitations that remain. First, MS, BA, display-based assays, and targeted presentation assays measure related but non-identical biological endpoints. Anchoring to MS is useful for retaining presentation knowledge. However, when updating with assay-specific knowledge, the true biological endpoint can become more difficult to characterize. Experimental results with paired outputs from specific assays and MS across peptide domains can provide a more thorough validation of how models updated with PepCL adopt new information in the context of natural peptide-MHC presentation. Second, deep peptide-MHC models are also sensitive to implementation choices, stochastic initialization, dataset splits, sampling schemes, negative construction, and hyperparameter settings. We performed all of our tests with controlled parameters, standard seeding, and minimal variation of hyperparameters to define a strong default baseline. We also interpret exact benchmark values cautiously and focus on patterns that recur across controlled synthetic tests, multiple assay updates, matched MS evaluations, and allele holdout settings.

Overall, PepCL reframes peptide-MHC prediction as a continual learning process rather than a one-time training task, shifting the field from a static paradigm toward an updatable framework for integrating new experimental evidence as it becomes available. This motivates further benchmarking and integration of the many non-MS assays that are often excluded from peptide-MHC model training despite providing complementary information. More broadly, our results suggest that progress will increasingly depend not only on improved model architectures, but also on frameworks that provide a practical path for expanding predictor coverage as new datasets and assay modalities emerge.

## Methods

### Data collection and processing

MS datasets were collected from prior studies and databases, including IEDB, MHC Motif Atlas, NetMHCpan4.1, MHCFlurry2.0, HLAthena, and HLApollo^8,9,10,16,56,57^. We filtered to mono-allelic peptides ranging from 8 to 14 residues and removed all flanking sequences and duplicate peptide-allele pairs.

Yeast display data were collected from Holec et al., 2025^19^. We used pathogen_library_ranks.xlsx to generate binder and non-binder distributions. EpiScan datasets were collected from the source data of Bruno et al., 2023^14^. Specifically, we used SD2, SD3, and SD6 for the random 9-mer (R9), SARS-CoV-2, and retraining library screens, respectively. For the R9 library, we used the SPP KO binders for HLA-A*02:01. We collected datasets for ESCAPE-seq from the source data of Shi et al., 2025^20^. The ESCAPE-seq data E-score values for the paired mutation library were thresholded at the recommended value of 3.8; peptides with scores greater than or equal to 3.8 are labeled as binders/positives, while all remaining peptides are labeled as non-binders/negatives. We additionally removed peptides containing “X” and data for HLA-A*68:01, as these peptides contained largely static E-score values. The reported neoantigen set from ESCAPE-seq was collected from Supplementary Table 7; the other peptide-allele pairs noted in the table were also expanded and included in the set. The IEDB and SARS-CoV-2 sets from ESCAPE-seq were thresholded at the recommended E-score value of 3.2. We collected binding affinity data from Reynisson et al., 2020^8^. The provided log50k transformed values were thresholded at 0.426 (values greater than or equal to 0.426 were labeled as binders, and all remaining peptides were labeled as non-binders).

We collected three MS pathogen-related datasets for SARS-CoV-2, Mycobacterium tuberculosis, and Zika from Weingarten-Gabbay et al., 2021, Leddy et al., 2023, and Sherwood et al., 2025, respectively^58,59,60^. Lastly, we collected cancer-related peptides from Ligand.MHC Atlas^61^.

We performed cleaning at both the peptide-allele and peptide levels, depending on the specific dataset distribution and tasks. Specifically, we removed all peptide-allele overlaps between MS and yeast display, EpiScan random 9-mer, EpiScan SARS-CoV-2, and BA datasets from the MS set. Peptide-allele overlaps between the MS and EpiScan retraining library were removed from the EpiScan retraining library due to the large number of binders and non-binders. For ESCAPE-seq and the MS pathogen-related datasets, we performed cleaning at the peptide level, removing overlapping peptides from the MS set. This was done to allow for more robust evaluation of rankability and deconvolution in downstream tasks.

The MHC pseudosequences were derived from Nielsen et al., 2007, and are all of length 34^62^. Swiss-Prot and human proteome data for generating the background distributions for ranking, synthetic negatives for training, and feature interpretability were collected from UniProt^63^.

The amino acid primitive and physicochemical features for the MHCPrime feature embeddings were collected from the peptides.py library. The features are split into two groups: primitive features and descriptor families. The primitive features include side-chain volume, isoelectric point, polarity, hydrophobicity, secondary-structure propensities, surface accessibility, turn propensity, and Miyazawa-Jernigan energy, and are standardized directly. The descriptor families included AF, BLOSUM, PP, F, KF, E, PD, PRIN, ProtFP, VHSE, and Z descriptors. Each descriptor family was standardized and reduced independently by PCA, retaining up to five principal components per family before final standardization. This yielded a 57-dimensional feature vector for each amino acid.

### Dataset distribution statistics and comparisons

To compare the sequence composition of non-MS assay binders with MS-derived ligands, we computed PWM divergence within matched allele-peptide-length groups. For each dataset, we retained binder peptides only and grouped peptides by allele and peptide length. Within each group, a PWM was computed from the observed amino-acid frequencies using a pseudocount of 1.0. Non-MS datasets were compared only in allele-length groups also present in the MS reference. For each matched group, the non-MS PWM was compared with the corresponding MS PWM using JSD, normalized by peptide length. The plotted values represent the resulting per-group divergences for each dataset category. As an MS reference, we estimated within-MS divergence by repeatedly splitting each MS allele-length group. In each of 10 iterations, 5% of peptides from the group were randomly sampled to form a query PWM, and this PWM was compared with a PWM computed from the remaining MS peptides. These within-MS values provide the reference distribution for the divergence expected between samples drawn from the same MS ligand repertoire.

In order to compute the percentage of binders mapping to the human proteome, we isolated all unique binders from each dataset and mapped them directly to the canonical human proteome (excluding isoforms). We then computed the fraction of peptides that mapped relative to all binders.

Cysteine enrichment was computed for each allele and dataset. We calculated the fraction of binder peptides containing at least one cysteine and the corresponding fraction among non-binder/background peptides. We then calculated the binder to background cysteine enrichment within each dataset and compared the non-MS enrichment ratio with the matched MS enrichment ratio for the same allele. Values are reported as log_2_ fold changes, so positive values indicate stronger binder cysteine enrichment in the non-MS dataset than in MS and negative values indicate weaker cysteine enrichment relative to MS.

Hydrophobicity shifts were computed using GRAVY scores based on the Kyte-Doolittle hydropathy scale^64^. For each peptide, the GRAVY score was calculated as the mean hydropathy value across its amino acids. We computed the mean GRAVY score of binder peptides for each allele in each dataset. The per-allele hydrophobicity shift was defined as the mean GRAVY score of non-MS binders minus the mean GRAVY score of matched MS binders for the same allele. Positive values indicate a hydrophobic shift relative to MS ligands, whereas negative values indicate a more hydrophilic shift.

### Train and test sets for training and evaluating MHCPrime

The MS test set used throughout this study was generated by taking 10% of binders from each allele, requiring a minimum of 5 and a maximum of 100 binders for each allele-length group. Synthetic negatives were sampled from the human proteome per allele-length group at a 1:99 positive to negative ratio for the test set. The remaining MS data were used to construct the train set. Synthetic negatives for the train set were sampled from the human proteome for each allele-length group at a 1:50 positive to negative ratio. Any peptide-allele or peptide overlaps with the collected datasets were appropriately excluded during the generation of synthetic negatives.

The yeast display, EpiScan random 9-mer, and EpiScan retraining library test sets were generated by sampling 100 positives per allele and sampling negatives at a 1:99 positive to negative ratio. Lastly, the BA and ESCAPE-seq test sets were generated by sampling the max possible number of positives and negatives to support a 1:99 positive to negative ratio. The BA and ESCAPE-seq ranking test sets for computing Spearman correlations were constructed with the max possible number of positives and negatives to support a 1:1 positive to negative ratio. All test sets were constructed to prohibit peptide-allele overlaps with the MS train set.

### MHCPrime architecture

The MHCPrime architecture follows a traditional dual-channel encoder, which is adapted from prior work on transformer-based peptide-MHC models. The two input channels accept a peptide sequence (lengths 8 to 14) that is right-padded with optional N- and C-flanking residues (max length 10; right-padded). The MHC channel accepts a pseudosequence of length 34.

Peptides and MHC pseudosequences are both tokenized and then fed into their respective channels to first produce the learned and feature embeddings. We use a total model embedding dimension of 480; 423 dimensions are learned while the remaining 57 dimensions are for the processed feature embeddings. To generate the merged learned and feature embeddings, we first initialize two separate embedding matrices for the peptide and MHC channels and generate the learned token embeddings for the peptide or allele pair.

In parallel, we also generate feature embeddings from the processed feature matrix, which maps each amino acid to a feature vector of length 57. We then use a linear projection shared across the peptide and MHC channels to map the feature matrix for each sequence into feature embedding space. This ensures that a given amino-acid token is mapped to a common projected biochemical feature representation before allowing for more specific weighting. We then perform channel-specific, context-aware gating to modulate the contribution of the feature embeddings. Specifically, per channel, we first add learned positional embeddings to the learned token embeddings and perform attention pooling over valid, non-padding positions to generate a context vector^65^. The context vector is then passed through a two-layer MLP to generate a feature gate. The feature gate is applied elementwise to the projected feature matrix to generate the final feature embeddings. This design allows the model to adjust the contribution of amino-acid feature dimensions as a function of the peptide or MHC sequence context. We then concatenate the respective feature embeddings to the learned token embeddings and add sinusoidal positional encodings along with segment embeddings. The segment encodings are added as a constant and are intended to distinguish the peptide (index 0) versus MHC (index 1) representations. This yields the full sequence-specific embeddings per channel.

The peptide and MHC embeddings then go through separate transformer/conformer encoder layers to output the final representations. Specifically, we use two Conformer blocks followed by two Transformer encoder layers in the peptide channel and a single Transformer encoder layer in the MHC channel. Transformer encoder layers are initialized with 16 attention heads and GELU activations in the feedforward layers. The Conformer block used in the peptide channel is adapted from Gulati et al., 2020, and is designed to extract both global and local dependencies^66^. We initialize the Conformer blocks with 16 attention heads and a convolution kernel size of 5.

The final peptide and MHC output representations from their respective channels are then concatenated, and a BOS token is added to the beginning. This concatenated representation is run through three more Transformer encoder layers and a single Conformer block with a convolution kernel size of 7. In addition, we mask the BOS token from the convolutional layers to avoid any proximity biases. Lastly, the BOS token is extracted from the concatenated output and passed through a three-layer feedforward neural network (FFNN; dimensions: 480 (BOS) → 240 → 120 → 1 (logit)) to produce a single output score. The FFNN uses a sigmoid GLU-style activation function^67^. The final model contained ∼25 million parameters.

### MHCPrime training

MHCPrime was trained on a single NVIDIA H100 GPU for 120 epochs using automatic mixed precision, the AdamW optimizer with a learning rate of 2e-4, default weight decay, a linear warmup scheduler, and a batch size of 3,072^68^. We use the strategy employed in NNAlign_MA and HLApollo for sampling peptide-allele pairs for training, where we sampled an equal number of positives and negatives per epoch (200k+/200k−), and negatives were sampled from a much larger pool to allow for negative cycling per epoch^10,69^. Since positive and negative allele-length compositions were the same in their respective label groups, this allows for balanced gradients. Because our downstream tasks largely involve recovery of positives among many negatives, we used a differentiable AP surrogate to align the training objective with the ranking metric of interest. Specifically, we use a LogSmoothAP ranking loss adapted from the Smooth-AP loss described by Brown et al., 2020^70^. Specifically, given model scores *s_i_* and binary labels*y_i_*, the hard pairwise indicator, 1(s_j_ >s_i_), is replaced with a sigmoid-based soft indicator function:

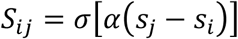

This yields a soft estimate of whether sample *j* is ranked above sample *i*. We use alpha (*α*) to control the strength of the logits and use a default value of 10.0. For each sample, soft precision can be computed as the ratio of soft positive count to the total soft sample count:

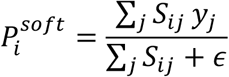

The batchwise smooth-AP is then computed by averaging these values over all positive examples:

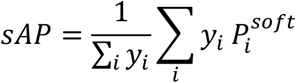

We minimized the negative log transform of the smooth-AP loss:

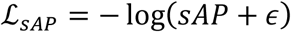

This form preserves the optimum of the smooth-AP while assigning a greater penalty to low smooth-AP batches and boundary samples.

### MHCPrime and external model benchmarking

We benchmarked MHCPrime and several published models across collected datasets, including HLApollo, DeepAttentionPan, TransPHLA, BigMHC EL, MUNIS, CapsNet-MHC IEDB and Anthem, MixMHCPred3.0, MHCFlurry2.0, NetMHCpan4.1 EL and BA, and NetMHCpan4.2 neoepitope (NE) and pathogen (PS) models^8,9,10,11,26,34,41,71,72,73^.

Full-length MHC protein sequences for alleles absent from the repositories of DeepAttentionPan, TransPHLA, MUNIS, and CapsNet-MHC IEDB were retrieved from IPD-IMGT/HLA and then formatted according to each model’s specifications. DeepAttentionPan and CapsNet-MHC IEDB output IC50 values which were then inverted. We used AP as our primary metric to evaluate performance on the MS and non-MS test sets. To summarize values, we use mAP across specified groups. For the MS test set, we additionally computed the area under the receiver operating characteristic curve (AUC), the partial area under the receiver operating characteristic curve up to a false positive rate of 0.1 (AUC0.1), and positive predictive value at n (PPVn). AUC summarizes classification performance across all thresholds by measuring the receiver operating characteristic curve, which plots true positive rate against false positive rate. AUC0.1 evaluates only the low false positive rate portion of the ROC curve, up to a false positive rate of 0.1. PPVn measures the fraction of true binders among the top n scoring peptides, with n defined as the number of positives in the evaluated set.

### PepCL method

PepCL begins by first initializing two copies of the provided model. One copy is used as the trainable model that is updated on incoming data while also receiving replay information if enabled. When drift regularization is also enabled, the second copy is used to penalize score differences from the original model score distribution. Training data is organized into an MS replay set and a new data set. The replay set contains samples from the original MS pretraining task, while the new data set contains the new assay or data source being integrated. We sample the sets separately because they can differ substantially in allele coverage, allele-specific sample counts, and positive to negative ratios. New data may also contain alleles that are absent, rare, or unevenly represented in the replay set, therefore, forcing the two sets to have matched allele distributions could oversample smaller allele subsets or discard useful new data structures. Instead, PepCL balances each set independently, while the relative contribution of replay and drift regularization is controlled by the batching and tunable lambda parameter.

Within each set, PepCL uses allele-aware sampling. For allele *a* in set *d*, let *P_d,a_* denote the number of positive samples for that allele. We compute a set-specific effective number of represented alleles using the standard effective sample size formula^74,75^:

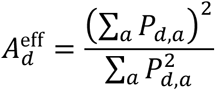

This value equals the number of represented alleles when positives are evenly distributed across alleles and becomes smaller when positives are concentrated in only a subset of alleles. We use this value to determine how strongly to balance allele sampling. Specifically, if *A_d_* is the number of alleles with at least one positive sample, we define:

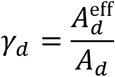

The allele sampling weight is then defined as a mixture of uniform allele sampling and count-proportional sampling:

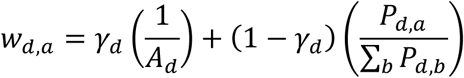

This weighted approach avoids high-count alleles from dominating training or repeated oversampling of low-count alleles under count-proportional or uniform sampling, respectively. By using the effective number of represented alleles, PepCL adjusts the degree of balancing according to the observed imbalance within each set. For each set, the target number of positive examples per sampling cycle is determined from the set batch size *B* and the negative to positive ratio *r_d_* :

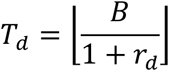

The maximum number of positives for an allele *a* is given by:

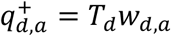

This is then converted into integer per allele positive counts and capped by the number of available positives for that allele. After positives are selected, negatives are sampled from the same allele at a 1:1 negative to positive ratio, subject to availability. Alleles without available negatives are still retained and contribute positive samples only.

During each training step, PepCL draws one batch from each set. In the standard setting, replay and new data batches are passed through the updated model copy separately, producing separate replay and new data losses whose gradients are accumulated before a single optimizer update. We use the same LogSmoothAP loss for both sets of data, staying consistent with how the base model was trained. Alternatively, for new data with continuous values, PepCL retains the LogSmoothAP classification objective on the MS replay set but replaces the new data classification loss with a differentiable soft rank loss. We use this instead of MSE to prioritize ordering over absolute magnitude, which could compete with the model logit scale. Exact rank operations are non-differentiable, so this loss was implemented using a smooth pairwise approximation to ranks, following the broader differentiable sorting and ranking literature rather than being directly adapted from a single existing loss^76^. Prior work has shown that hard sorting and ranking operations can be replaced with differentiable relaxations, including differentiable Spearman-style objectives; this is also analogous in spirit to smoothed rank-metric objectives such as Smooth-AP, although our continuous value objective optimizes rank correlation rather than AP^77,78^. Given model scores *s_i_* and continuous targets *z_i_* in a new data batch of size *B*, we compute soft ranks using a pairwise sigmoid form:

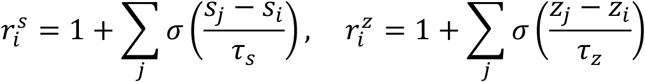

where *τ_s_* and *τ_z_* are temperature values used to scale the model scores and target values, respectively. We next center the predicted and target soft rank vectors:

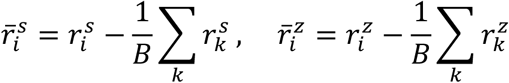

The normalized correlation is then computed as:

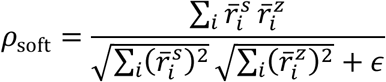

The final continuous loss is:

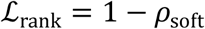

By default, we use *τ_s_* = 0.01 for the model scores and *τ_z_* = 10^−4^ for the target values, with *ε* = 10^−8^. These temperatures control the sharpness of the rank approximation where smaller values more closely approximate hard ranks and larger values produce smoother pairwise comparisons. We kept the target temperature small so that the target ranks closely reflect the empirical ordering of the assay-specific continuous values, while the prediction temperature can be adjusted to control gradient smoothness. For datasets with different target scales or measurement noise, these temperatures can be varied to test more or less granular rank sensitivity.

When drift regularization is enabled, the replay set is additionally passed through the frozen model copy and loss is computed as the mean squared error (MSE) between the replay logits through the updated and frozen model copies. We use MSE here because the model scores are unbounded rather than being calibrated probability distributions. Although a Kullback-Leibler divergence loss could be defined after converting logits to sigmoid probabilities, this conversion can compress large magnitude logits into near binary values and reduce sensitivity to the frozen model’s score margins. MSE therefore provides a simple way to preserve the frozen model’s replay score distribution when updating with new data.

Lastly, with replay and drift regularization enabled, each training step requires a forward pass of the updated model on the replay batch, a forward pass of the updated model on the new data batch, and a forward pass of the frozen model on the replay batch. Because the frozen model does not change during training, frozen model replay logits can optionally be cached, eliminating the repeated frozen model forward pass and reducing the online computation to the two updated model forward passes.

### Log odds computation

We computed an allele- and length-specific PWM log odds score from a given set of positive ligands. For each allele-length group, amino acid frequencies were estimated at each peptide position using the given positive ligands as the reference set. Background amino acid frequencies were estimated separately for each peptide length from the corresponding background peptide pool. Reference PWM frequencies were regularized toward the background distribution using a pseudocount of 50.0, and background frequencies were smoothed using a pseudocount of 1.0. The log odds contribution for each peptide position was then computed as the log ratio between the allele-specific PWM probability and the background amino acid probability, and the final peptide score was the sum of these positionwise log odds values across the peptide.

### Synthetic MS in-distribution analysis

The train set was split into 10 sets containing 10% of the data each. The first 10% contained rarer alleles with lower than 50 binders. We trained a series of base models with increasing increments of data and then updated the model with the next 10% split using a lambda value of 0 and 1,000 steps. The log odds were computed on the test set using the binders from each split. The mean AP (mAP) per model was computed by averaging the AP per allele. The AP differences between models were computed by taking the difference between the current pretrained model per allele AP and the model updated on the next split of data. We then averaged the per allele AP differences across all nine splits.

### Allele-specific cluster holdout updating and analysis

For six alleles, HLA-A*02:01, HLA-A*03:01, HLA-B*27:05, HLA-B*40:02, HLA-C*05:01, and HLA-C*12:02, we first embedded all the MS 9-mer binders using the processed feature tables. This generated feature vectors representing each peptide. We constructed a weighted k-nearest-neighbor graph using cosine similarity with k = 30, retaining only mutual nearest-neighbor edges. Leiden clustering was then performed on the weighted graph using a resolution parameter of 1.2. Clusters containing fewer than 50 peptides were grouped into a single small cluster^79,80^.

For each allele-specific cluster, we held out 10% of binders with a maximum of 150 peptides for the test set, along with paired synthetic negatives at a 1:99 positive to negative ratio. For HLA-A*02:01, we used 150 binders from each cluster as it was the most abundant allele. We additionally sampled cluster-specific negatives at a 1:99 positive to negative ratio from the human proteome by scoring remaining sequences with cluster-specific binder log odds and selecting the highest scoring background sequences per cluster.

For each allele-specific cluster, we trained a model with the cluster held out, using only the 9-mer training data from other clusters for the target allele while keeping the remaining MS train data constant. We additionally trained a set of models with the entire allele held out and full models in which we used all the 9-mers from the target allele across the resolved clusters. For each cluster held-out model, we updated with cluster-specific binders paired against synthetic negatives. The replay set consisted of data from all remaining clusters along with the same synthetic negatives. We used 200 steps and a lambda value of 1e-4 for PepCL. For all models, including the updated ones, we computed the AP of all clusters and then averaged the AP of all non-target clusters to derive a prior mAP. Therefore, each model was used to compute a target cluster AP and prior mAP, including the full models and allele held-out models for consistency.

To test how the update cluster’s distance to other clusters was related to the extent of forgetting on the remaining clusters, we compared cluster-specific motif divergence with the change in held-out performance after cluster-specific updating. For each of six alleles, we evaluated models updated on individual peptide clusters and computed AP separately for each held-out cluster. For a model updated on a given target cluster, held-out mAP was calculated as the mean AP across all non-target clusters. The same held-out mAP was computed for the corresponding base MHCPrime scores, and ΔmAP was defined as the updated model held-out mAP minus the base held-out mAP. To quantify how distinct each target cluster was from the remaining ligand repertoire, cluster PWMs were computed from allele-specific MS 9-mer training ligands using a pseudocount of 0.5, and the mean JSD from the target cluster to all other clusters was calculated and min-max normalized. We then fit an ordinary least-squares linear regression between normalized cluster-to-rest divergence and ΔmAP across all allele-cluster pairs. The shaded band shows the 95% confidence interval for the fitted mean regression line, and Pearson and Spearman correlations were also computed.

### Unseen allele analysis

We used our MS train set and first identified alleles that contained less than 10,000 but more than 1,000 binders. This provided a set of medium prevalence alleles from which we generated 10 sets of 10 alleles each to hold out from during training. For each allele set, we trained our base model with all data while holding out the target allele set. We then updated each model with the target allele set while using the remaining data as the replay set for PepCL. We used default parameters for all updating experiments (200 steps; λ = 1e-5). For each updated model with its respective allele split, we computed the mAP of the pretraining and held-out alleles separately.

### Yeast display updating

We merged the MS and yeast display train sets after removing peptide-allele overlaps with the test sets specified above. We then updated the base MHCPrime model using 200 steps with either no replay (FT) or replay with a lambda value of 1e-5. For PepCL, the replay set was matched to the allele and length distribution of the available yeast display data, yielding 16 alleles containing 9-mers only. For all models, we computed the allele-specific AP of the yeast display test set and the matched allele-length MS test set.

To quantify the per-allele trade-off between the improvement on the yeast display set and preservation of MS performance, we computed a net benefit score. For each allele, we first measured the AP gain on the yeast display test set relative to the base MHCPrime model. We then computed the MS performance loss relative to the same base model. The net benefit was defined as the yeast display AP gain minus the MS AP loss. Positive values indicate that the gain on yeast display data exceeded the loss on matched MS data. Negative values indicate that MS loss was larger than the yeast display improvement.

For the mixture analysis, mixed test sets were then generated by varying the fraction of examples drawn from MS and yeast display in 10% increments, ranging from 100% yeast display to 100% MS. All test sets were held at a 1:99 positive to negative ratio, and the MS set was matched to the allele-length distribution of the yeast display set. At each mixture proportion, we performed subsampling 100 times. In each iteration, positives and negatives were sampled without replacement within allele and assay according to the specified mixture fraction. For each mixed test set, FT and PepCL mAP (per-allele mean AP) were compared against the base MHCPrime model on the same sampled examples, yielding a paired ΔmAP relative to the base model. The 95% bootstrap confidence intervals were computed from the 2.5th and 97.5th percentiles across iterations. Lastly, we performed allele deconvolution and ranking on the MS pathogen sets as specified in the ranking and recovery section.

### EpiScan updating

We first created mixed training sets for the EpiScan R9 and RT libraries by pairing the R9/RT+ with either R9− or R9/RT− samples. We then updated the base model with each dataset using FT and PepCL using the default parameters (200 steps; λ = 1e-5). For PepCL, we used a matched allele-length replay set. We gauged performance of all updated models by scoring mixtures of the R9 and RT test sets. We specifically sampled random mixtures of R9/RT+ paired against randomly subsampled R9− samples for the easy negative background test set. For the harder negative background test set containing RT−, we randomly sampled mixtures from R9/RT−. We maintained a 1:99 positive to negative ratio across all splits. Sampling was performed 100 times for each set, and the per allele AP was computed by averaging the AP values across iterations. The displayed mAP is computed by averaging the per allele AP values.

For the EpiScan SARS-CoV-2 set, we restricted to the matching allele-length groups and performed repeated subsampling (100 times) of the negatives to achieve a 1:99 positive to negative ratio. The per allele AP was computed by averaging the AP values across iterations, and the final mAP was computed by averaging the per allele AP values.

### Binding affinity updating

We resampled a test set with 5% of the binders per allele, requiring a minimum of 40 and maximum of 200, and negatives at a 1:1 positive to negative ratio. We used the remaining data for updating. This yielded 83 alleles for updating and 51 matched alleles for benchmarking. The base model was updated with the log50k BA values under a classification and continuous setting. For PepCL, the replay set contained matched allele-length samples and used the traditional LogSmoothAP classification loss for the replay set. We used default parameters for updating (200 steps; λ = 1e-5). We then computed the AP and Spearman correlation to the BA values per allele for all models for the BA and MS test sets.

### ESCAPE-seq updating

We first held out the reported neoantigens from ESCAPE-seq at the peptide level and updated on all remaining data. We specifically performed both classification and continuous value updating using the provided E-scores. We additionally updated under the classification pathway using additional synthetic negatives in the new data set (**Supplementary Fig. 8**). For PepCL, the replay set contained matched allele-length peptides from MS. We did not remove alleles from ESCAPE-seq that were not present in the MS training data as the reported neoantigens contained several peptides labeled with those alleles. Default parameters were used for updating (200 steps; λ = 1e-5). We scored the reported neoantigens and derived ranks as described above for the base and updated models. We additionally scored and ranked the Ligand.MHC Atlas test data to gauge recovery on an external cancer set. The Ligand.MHC Atlas test set was generated by removing all peptide-level overlaps with the MS and ESCAPE-seq update sets then matching allele and length distributions.

For the differential mutation calling test, we held out data corresponding to each gene or gene group, as some genes contained overlapping peptide sets. For the holdout set, we prevented any raw sequence overlaps with the train or new data. We additionally removed duplicate peptide-allele pairs that had mixed E-score signs from the train set; we specifically removed the negative peptide-allele pairs while keeping the positive scoring pairs in line with maintaining a positive preference. We performed FT and PepCL updating and then scored the gene holdout set. The replay set for PepCL used the same methodology for the reported neoantigen recovery task. In order to pair wild-type and mutant peptides for the differential mutation scoring analysis, we uniquely indexed the wild-type and mutant dataframes by the “name2”, “allele”, and “gene” columns, then matched pairs across dataframes while ensuring at least one edit was made between matched peptides. We computed score differences for all pairs by subtracting the wild-type peptide score from the mutant peptide score. The gain and escape mutations were labeled based on the E-score thresholds. Specifically, gain mutations are wild-type peptides that have E-score values below 3.8 and have paired mutant peptides with E-score values above 3.8. Escape mutations are the opposite. The AP was computed at the gene/gene group level as opposed to the allele level as each set contained a mixed set of alleles. The score differences were reversed to compute the per gene/gene group AP values for the escape mutations, since those pairs required identifying targets with lower values for the positive class (1) than the negative class (0).

### Allele transfer learning

We used both EpiScan R9− and R9/RT− models and the base MHCPrime model to score the remaining eight alleles in the EpiScan SARS-CoV-2 library. For 9-mers only, we then repeatedly sampled negatives at a 1:99 positive to negative ratio and computed allele-specific AP. The per-allele AP was computed by averaging allele-specific AP across 100 iterations. The difference from base was computed by taking the difference between each updated model’s allele-specific AP and the base model’s allele-specific AP.

For the yeast display, BA, and ESCAPE-seq datasets, we specifically held out each allele while updating on the remaining data. We used allele-length matched replay sets for PepCL and used 9-mers only for both update methods. We updated with default parameters (200 steps; λ = 1e-5). For each dataset, allele-specific AP was estimated by repeated stratified subsampling. For each allele, we sampled the maximum number of positive examples that could support a 1:99 positive to negative ratio, sampled the corresponding number of negatives, and computed AP. This was repeated 100 times per allele, and allele-specific AP was averaged across iterations.

### Validation of assay-specific properties for updated models

To measure enrichment in EpiScan-specific properties, we first sampled 100,000 peptides from the human proteome and scored them under the four available alleles for the EpiScan R9/RT− models and the base MHCPrime model. For all analyses, we retained the top 2,000 (2%) scoring peptides per allele. Cysteine enrichment was computed as the fraction of top-ranked peptides containing at least one cysteine. For each updated model, this fraction was compared with the corresponding fraction from the base MHCPrime top-ranked set for the same allele and reported as the log_2_ enrichment relative to base. Positive values indicate that the updated model ranked more cysteine-containing peptides among its top predictions than the base model, whereas negative values indicate reduced cysteine representation among top-ranked predictions. Hydrophobicity shifts were computed on the same top-ranked peptide sets. Each peptide was assigned a GRAVY score by averaging residue hydropathy values using the Kyte-Doolittle scale. For each allele and model, we calculated the mean GRAVY score among the top 2,000 ranked peptides. Updated models were then compared with the base MHCPrime model by subtracting the base model mean GRAVY score for the same allele. Positive values indicate that model updating shifted top-ranked predictions toward more hydrophobic peptides, whereas negative values indicate a shift toward less hydrophobic peptides.

For the penultimate proline enrichment and the C-terminal V/L shift, we also used the top 2,000 scoring peptides and computed these for HLA-A*02:01 and HLA-B*08:01. We specifically compared the updated models and the EpiScan log odds to the base MHCPrime model. We computed the penultimate proline enrichment in the same fashion as the cysteine-containing peptide enrichment, counting peptides with proline at position 8. For the C-terminal V/L shift, we counted the number of top-ranked peptides ending in valine and leucine and calculated the V/L ratio. Updated models and the EpiScan log odds score were then compared with the base MHCPrime model for the same allele and reported as log_2_ shifts in the V/L ratio. Positive values indicate that the scoring method shifted top-ranked predictions toward relatively more C-terminal valine compared with leucine, whereas negative values indicate a shift toward relatively more C-terminal leucine.

For the Y84A-related analyses, we randomly sampled and scored 100,000 peptides from Swiss-Prot for all 16 yeast display alleles using the base MHCPrime model and the updated models. For each allele and model, we retained the top 2,000 scoring 9-mer peptides and computed position-specific amino acid distributions across the selected peptides. Positional entropy was computed from these distributions. To compare FT and PepCL, we subtracted the PepCL positional entropy from the FT positional entropy for each allele and position; positive values indicate that FT selected a more compositionally diverse set of peptides at that position, whereas negative values indicate that FT selected a sharper positional distribution than PepCL. To quantify loss of composition relative to the base model, we also computed position-specific JSD between the amino acid distribution of the top 2,000 peptides selected by each updated model and the corresponding top 2,000 peptides selected by the base MHCPrime model for the same allele. We then subtracted the PepCL divergence from the FT divergence at each allele and position. Positive values indicate that FT deviated more strongly from the base model positional composition than PepCL, whereas values near zero indicate similar deviation from base.

### Ranking and recovery of peptide-MHC pairs

For rank-based evaluation settings, model scores were converted to global background percentile ranks, following the ranking strategy used in HLApollo. A background peptide set was generated from the human proteome and scored with the same model used to score the corresponding test peptides. Each test peptide was then assigned a percentile rank according to its position within this global background score distribution. Ranks were oriented such that higher values indicate a peptide scored above a larger fraction of the background set and therefore correspond to stronger predicted binding or presentation. The same procedure was applied to the base and updated models, ensuring that rank comparisons reflected model-specific score distributions against an identical background.

For recovery-related tasks on multi-allelic sets (MS pathogen sets), we computed the rank values for a given peptide paired against all possible alleles in the allotype and assign the peptide the allele with the maximum rank value. We computed the average rank using all possible pairings. For mono-allelic sets (ESCAPE-seq reported neoantigens and Ligand.MHC Atlas test set), we also compute the average rank and the percentage of binders that cross the 2% threshold (framed as greater than or equal to 98% for our inverse rank values).

### P-value computation

Statistical significance was assessed using paired Wilcoxon signed-rank tests. Comparisons were performed on matched groups, such as the same alleles, datasets, or evaluation splits, depending on the analysis.

### Code availability

The packaged versions of MHCPrime (https://github.com/ntranoslab/MHCPrime) and PepCL (https://github.com/ntranoslab/PepCL) are available on GitHub.

### Data availability

All MS and non-MS peptide-MHC datasets were obtained from publicly available sources described in the methods.

### Declaration of AI usage

During preparation of this manuscript and associated analysis code, the authors used ChatGPT (OpenAI), Claude (Anthropic), and Gemini (Google) to assist with language editing and code drafting. All AI generated code was reviewed, modified where necessary, tested on the relevant datasets, and executed by the authors. All AI generated text was reviewed and edited by the authors. The authors take full responsibility for the accuracy, reproducibility, interpretation, and integrity of the final manuscript, figures, and code.

## Acknowledgements

We thank Sohum Kothavade for helpful discussions and technical guidance during the development of the model architecture and continual learning framework.

## Competing Interests

V.N. is a scientific advisor for Tendel Therapies. P.M.B. is co-inventor of EpiScan PCT/US2020/062912. P.M.B. also serves on the scientific advisory board of and holds equity in Sift Biosciences. All other authors have no competing interests to declare.

---

**Supplementary Fig. 1:**
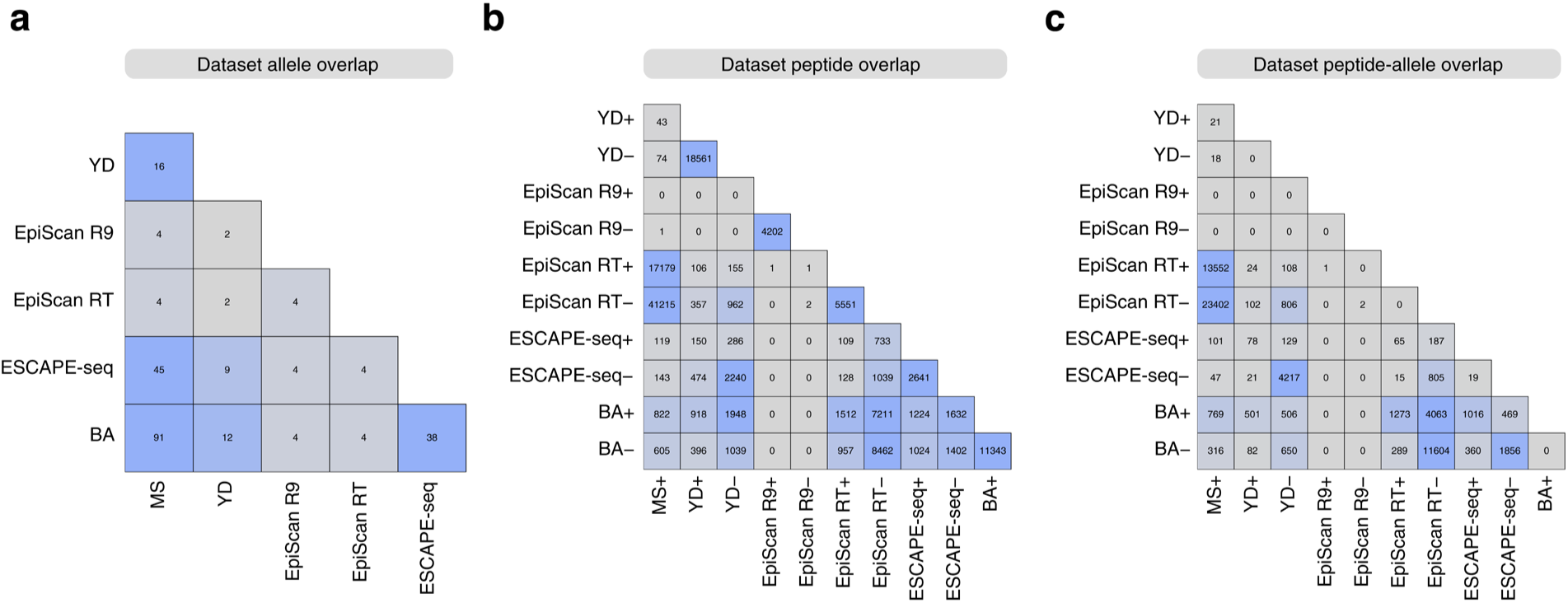
**Dataset overlaps before filtering**. **a)** Allele overlaps between the unfiltered datasets used in this study before cleaning and overlap removal. **b)** Peptide sequence overlaps between unfiltered datasets. **c)** Peptide-allele pair overlaps between unfiltered datasets. + and − symbols indicate whether the corresponding dataset contains binders and/or non-binders.

**Supplementary Fig. 2:**
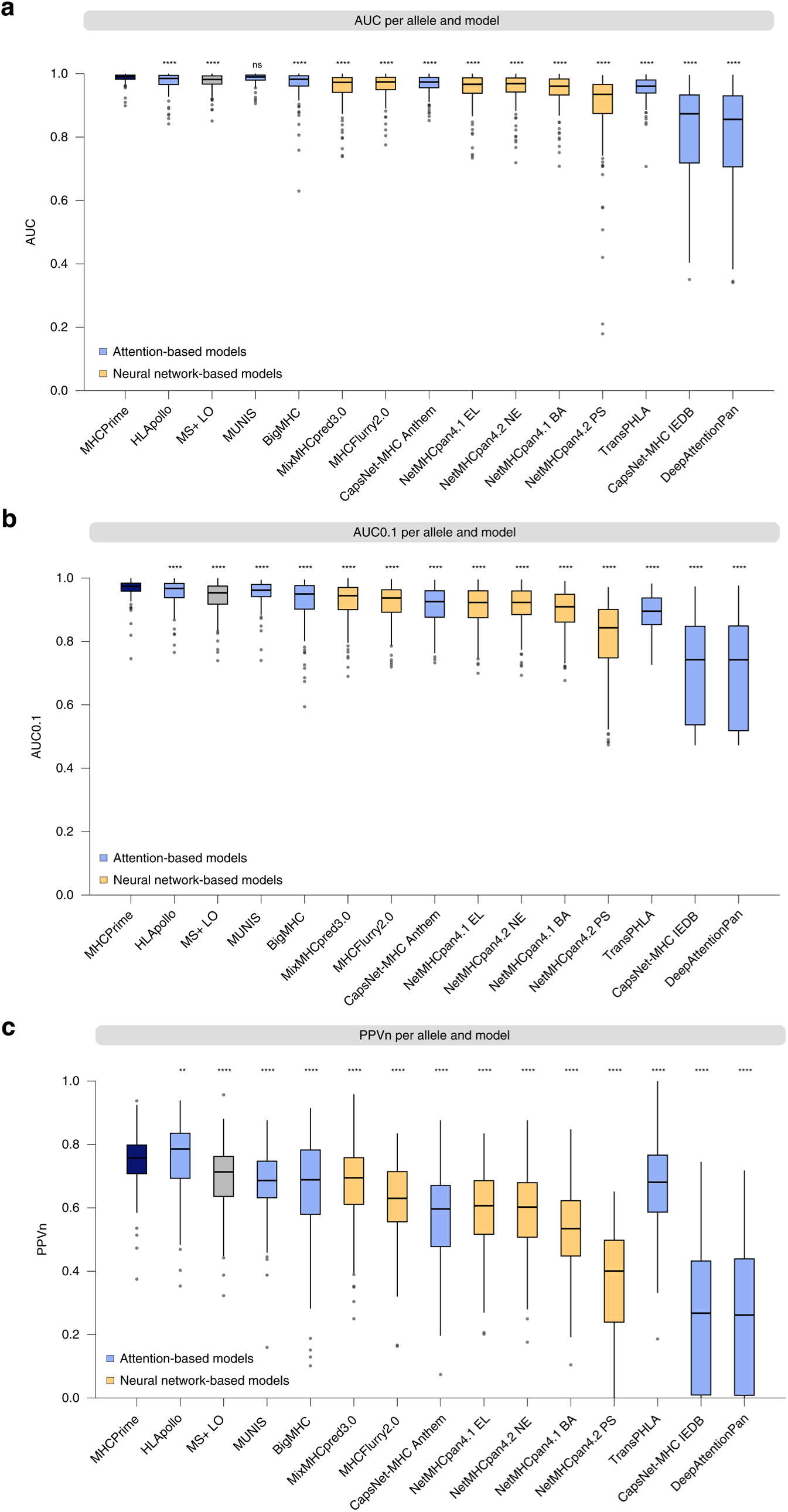
**Additional MS benchmark metrics**. **a)** Per-allele AUC on the held-out MS test set across benchmarked models. **b)** Per-allele AUC0.1 on the held-out MS test set across benchmarked models. **c)** Per-allele PPVn on the held-out MS test set across benchmarked models. All metrics were computed using the same MS test set and 1:99 positive to negative ratio used for the main AP benchmark. P-values were computed with paired Wilcoxon signed-rank tests relative to MHCPrime (significance labels indicate: ns, P > 0.05; *, P ≤ 0.05; **, P ≤ 0.01; ***, P ≤ 0.001; ****, P ≤ 0.0001). Colors denote attention-based and neural-network-based models.

**Supplementary Fig. 3:**
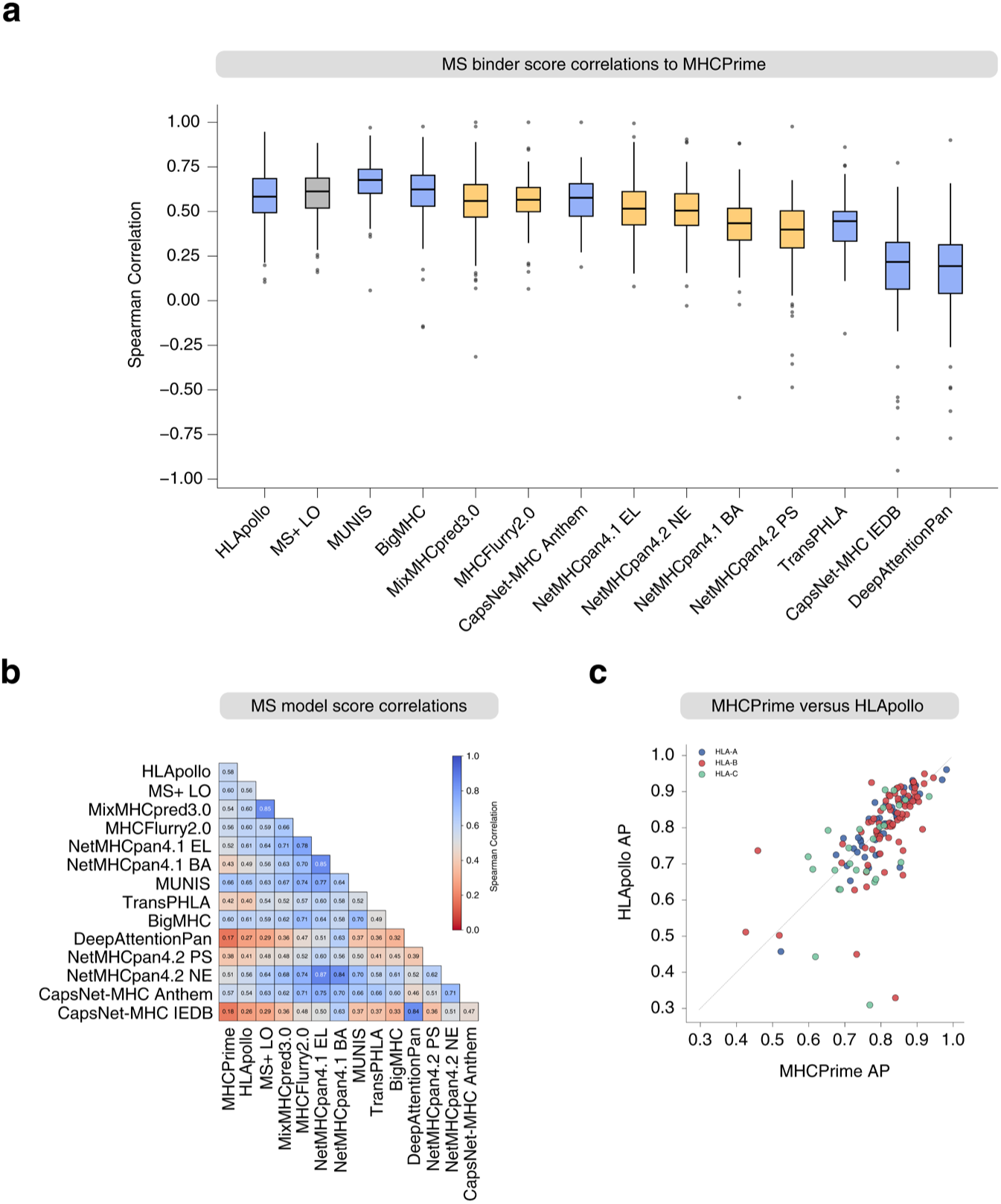
**Score concordance between MHCPrime and external predictors**. **a)** Per-allele Spearman correlation between each model and MHCPrime on MS positive test peptides, computed across 141 alleles. **b)** Average pairwise Spearman correlation between model scores on the MS test set. Pairwise correlations were computed within each allele using all MS test peptides, then averaged across alleles. **c)** Per-allele AP comparison between MHCPrime and HLApollo on the MS test set, with points colored by HLA locus.

**Supplementary Fig. 4:**
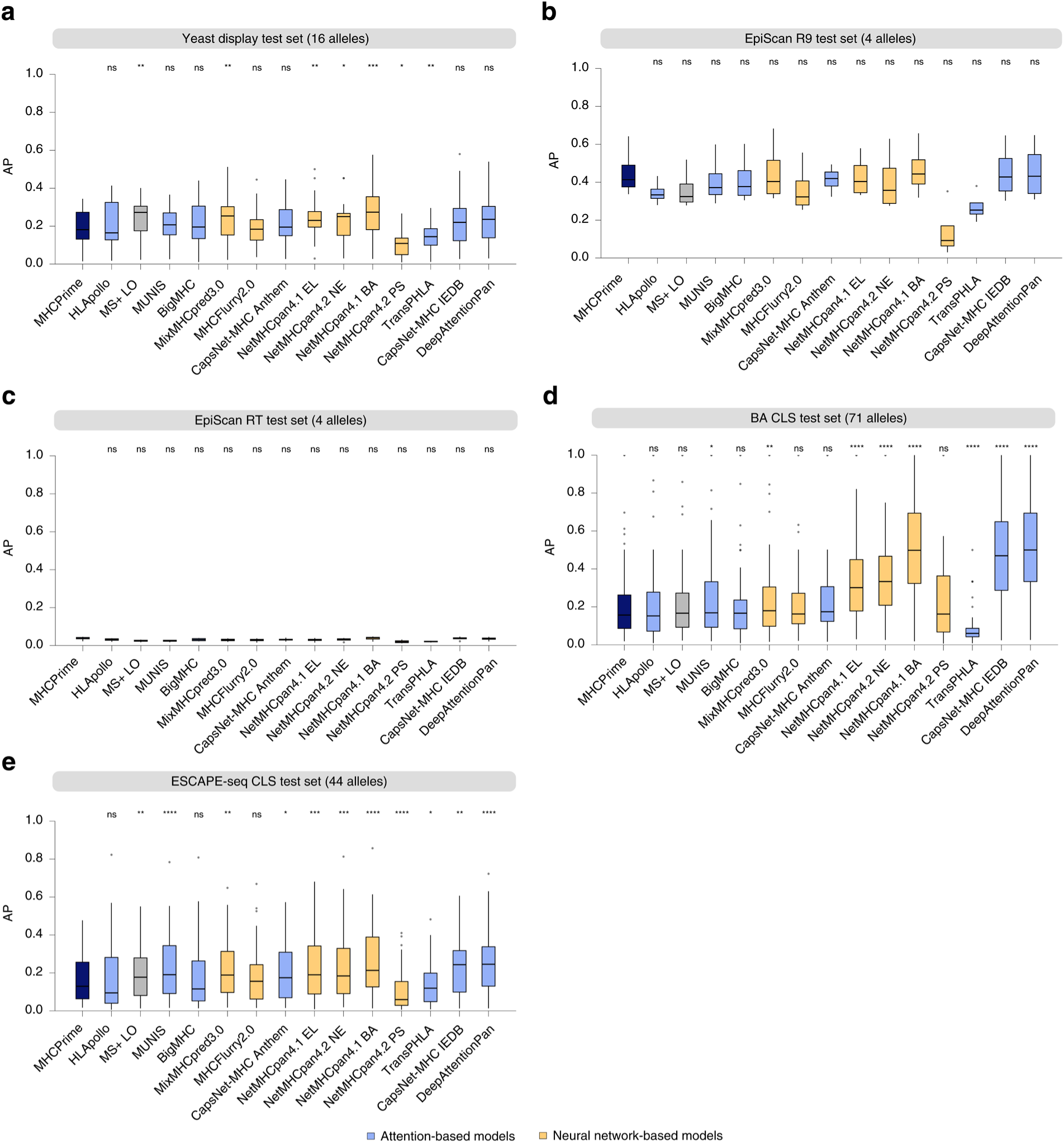
**Non-MS benchmark AP distributions**. **a-e)** Per-allele AP distributions for the non-MS datasets summarized in Fig. 2e. Each dataset was evaluated at a 1:99 positive to negative ratio using the available allele-length groups for that benchmark. P-values were computed with paired Wilcoxon signed-rank tests relative to MHCPrime within each dataset (significance labels indicate: ns, P > 0.05; *, P ≤ 0.05; **, P ≤ 0.01; ***, P ≤ 0.001; ****, P ≤ 0.0001). Colors denote attention-based and neural-network-based models.

**Supplementary Fig. 5:**
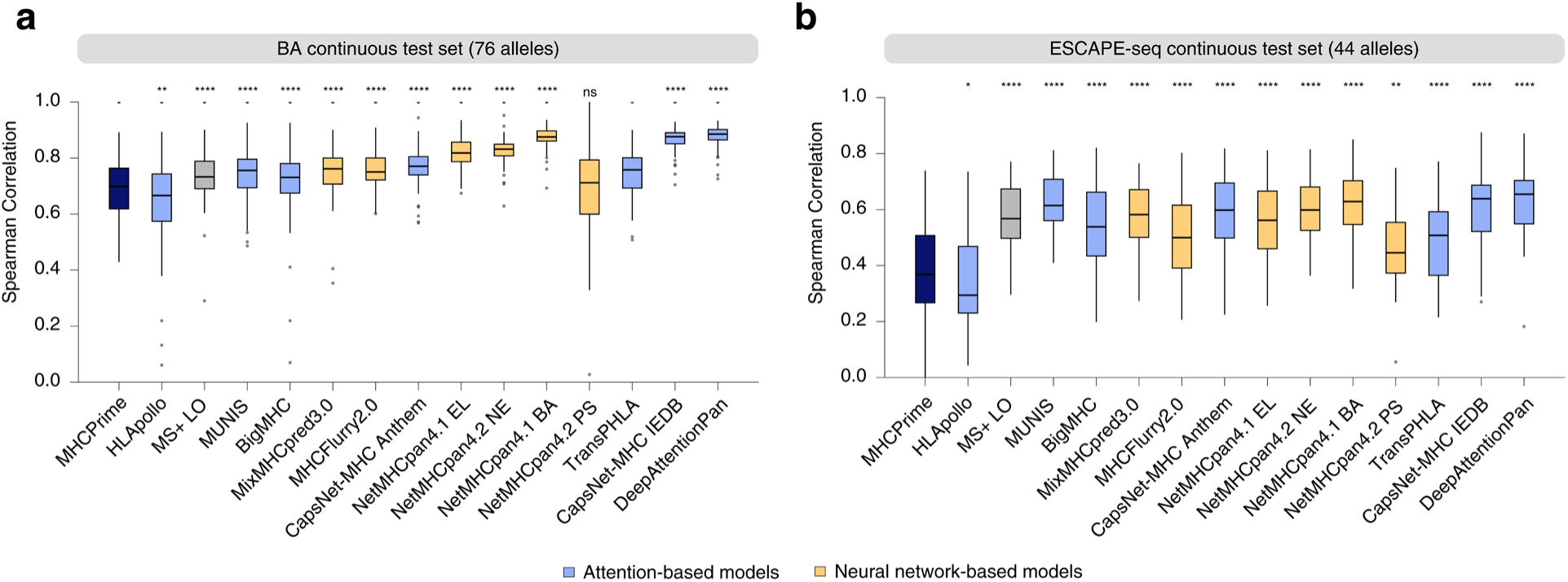
**Continuous-readout benchmark correlations**. **a)** Per-allele Spearman correlation between model scores and BA values. **b)** Per-allele Spearman correlation between model scores and ESCAPE-seq E-scores. BA and ESCAPE-seq test sets were evaluated at a 1:1 positive to negative ratio. P-values were computed with paired Wilcoxon signed-rank tests relative to MHCPrime (significance labels indicate: ns, P > 0.05; *, P ≤ 0.05; **, P ≤ 0.01; ***, P ≤ 0.001; ****, P ≤ 0.0001). Colors denote attention-based and neural-network-based models.

**Supplementary Fig. 6:**
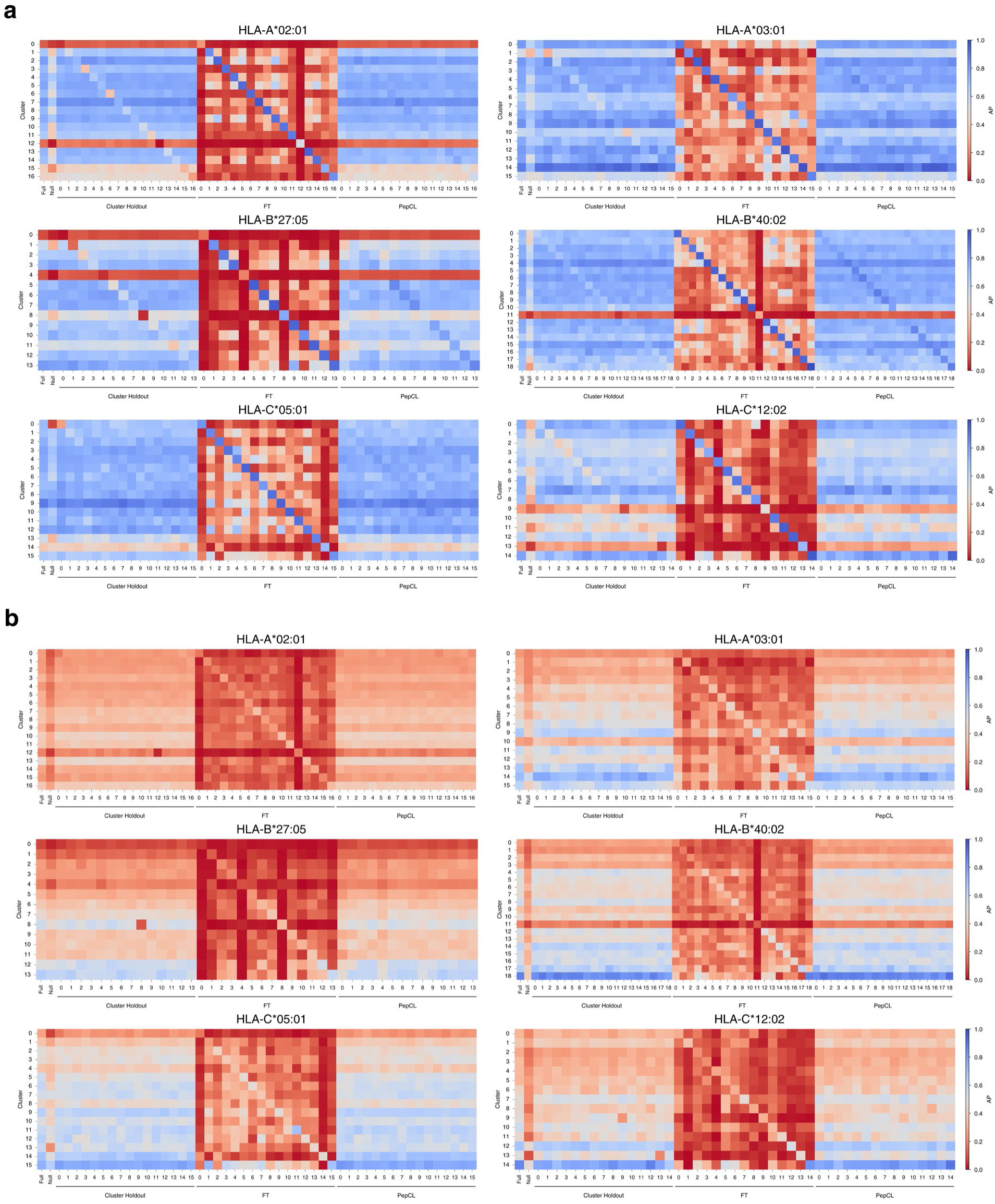
**Cluster-specific synthetic update performance**. **a)** Cluster-specific AP for held-out cluster updates evaluated with general negatives. **b)** Cluster-specific AP for the same held-out cluster updates evaluated with hard cluster-specific negatives. The x-axis indicates the target cluster. Full indicates the model trained with all clusters, Null indicates the model trained without the corresponding allele, Cluster Holdout indicates the model trained with the target cluster removed, and FT and PepCL indicate models updated with the originally held-out target cluster. Individual values represent AP for each cluster-specific evaluation.

**Supplementary Fig. 7:**
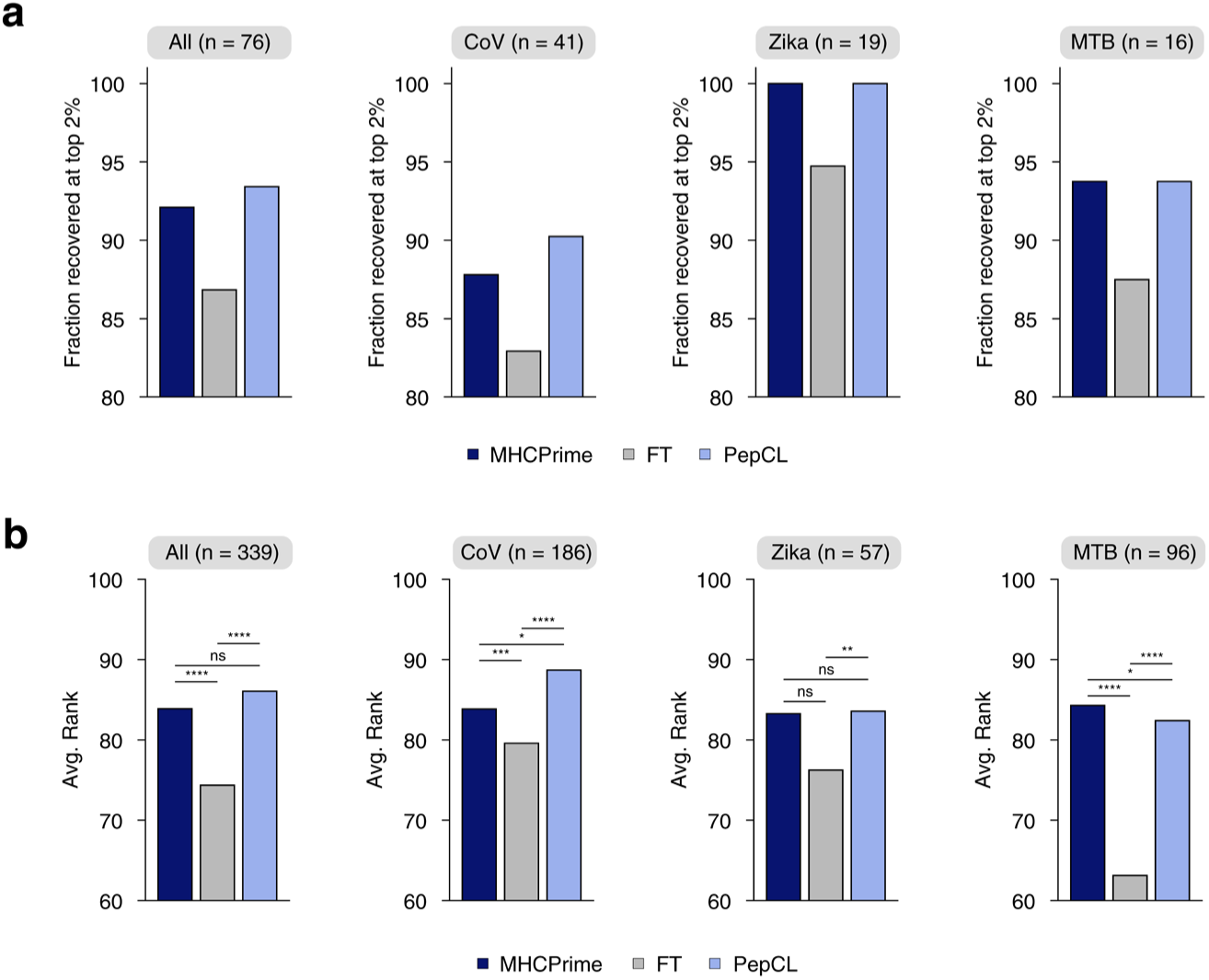
**Ranking and recovery of MS pathogen sets after yeast display updating**. **a)** Fraction of pathogen MS peptide groups recovered above the top 2% rank threshold after scoring all possible peptide-allele pairings within each sample or donor allotype. For each peptide group, the highest ranking compatible peptide-allele pairing was used to determine recovery. Results are shown for all pathogen ligands combined and separately for the indicated pathogen groups. **b)** Average rank across all compatible peptide-allele pairings for the same pathogen MS sets. P-values were computed with paired Wilcoxon signed-rank tests across peptide-allele pairs relative to MHCPrime (significance labels indicate: ns, P > 0.05; *, P ≤ 0.05; **, P ≤ 0.01; ***, P ≤ 0.001; ****, P ≤ 0.0001).

**Supplementary Fig. 8:**
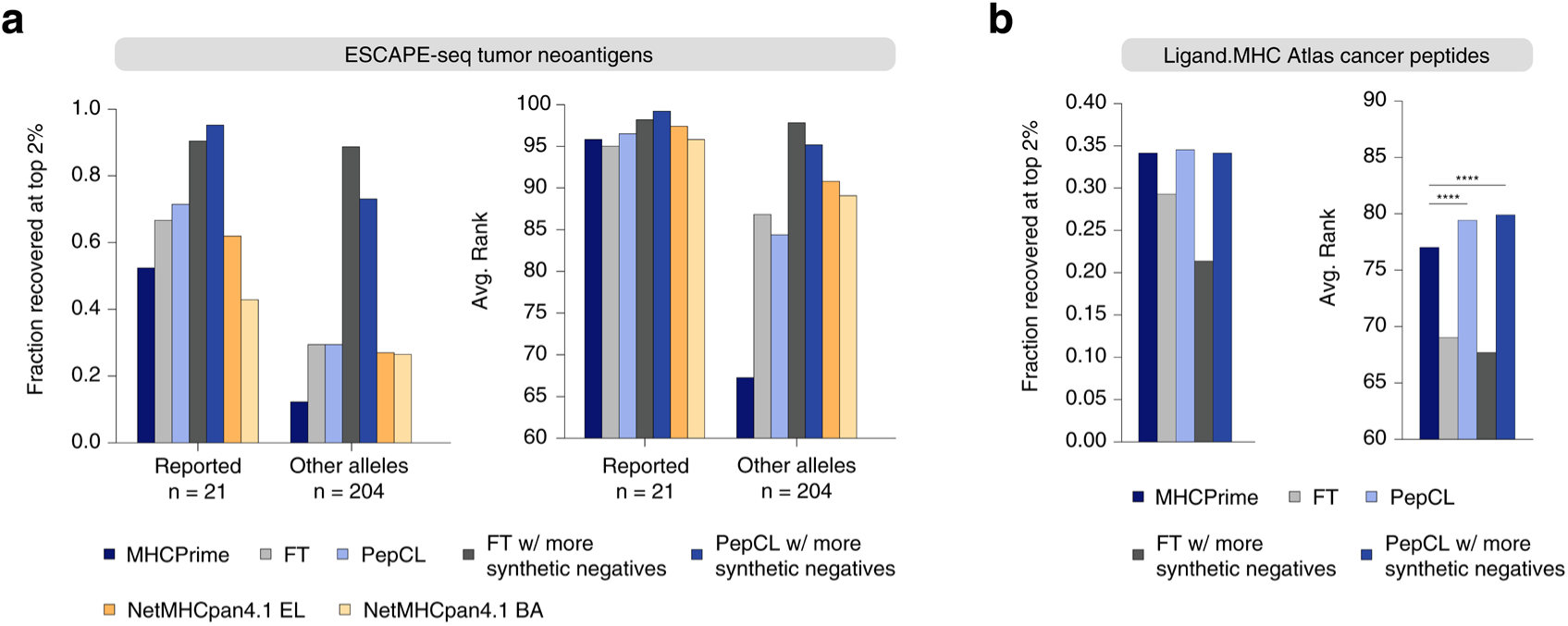
**Additional recovery benchmarks after ESCAPE-seq updating**. **a)** ESCAPE-seq tumor neoantigen recovery. **Left**, fraction of reported and alternative allele-paired peptide-allele pairs recovered above the top 2% rank threshold. **Right**, average rank for the same peptide-allele pairs. Darker model variants indicate updates performed with additional synthetic negatives sampled from matched MS allele-length distributions. **b)** Ligand.MHC Atlas recovery using monoallelic ligands with no raw peptide overlap with the update or training sets. **Left**, fraction recovered above the top 2% rank threshold. **Right**, average rank. All samples shown are mono-allelic and did not require deconvolution.

**Supplementary Fig. 9:**
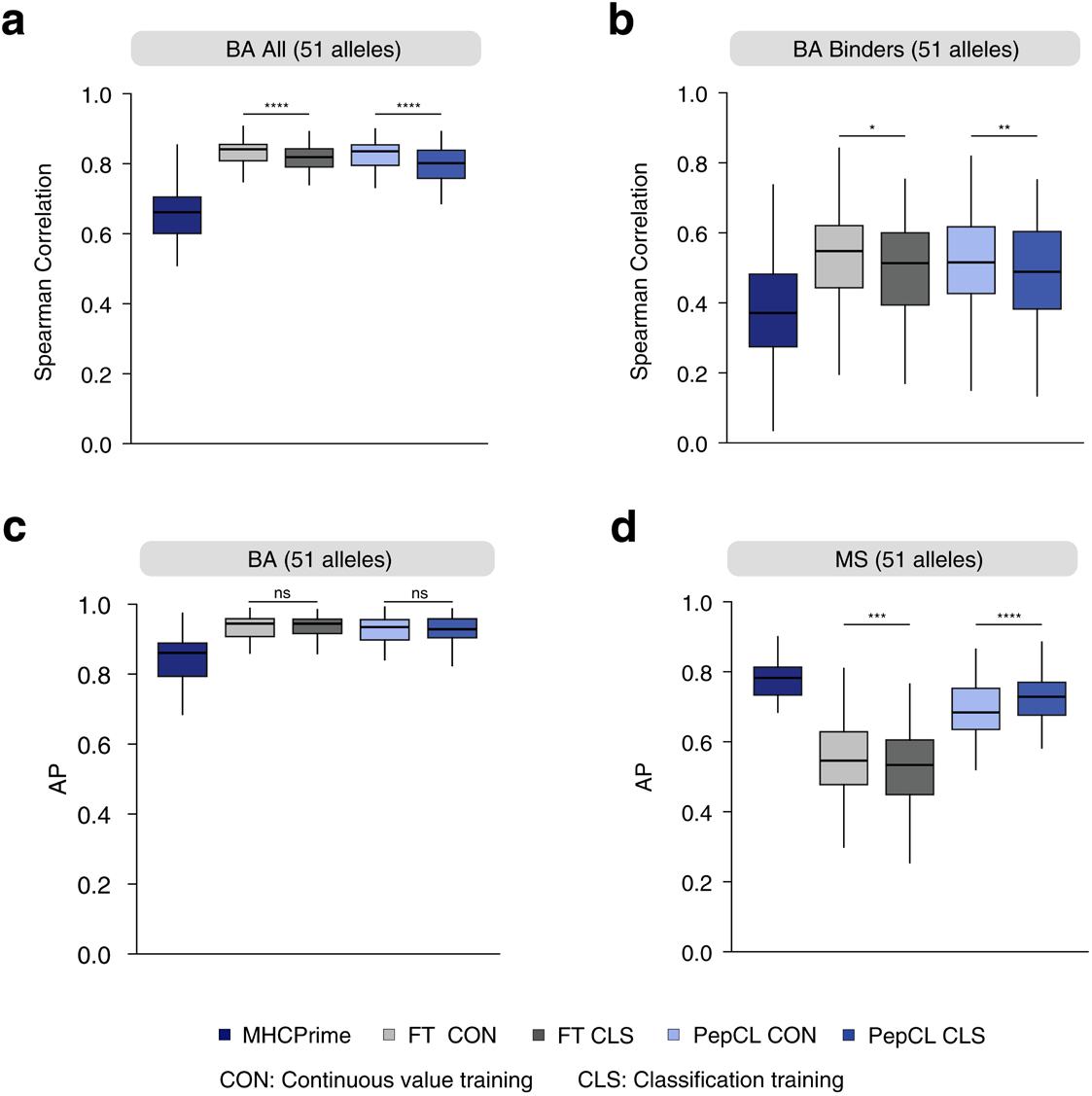
**Additional BA update benchmarks**. **a)** Per-allele Spearman correlation between model scores and BA values across the BA test set evaluated at a 1:1 positive to negative ratio. **b)** Per-allele Spearman correlation between model scores and BA values among BA binders only. **c)** Per-allele BA AP distributions. **d)** Matched MS AP after BA updating. Continuous-value models were trained with a continuous ranking objective, whereas classification models were trained using thresholded BA labels. P-values compare the indicated model groups (significance labels indicate: ns, P > 0.05; *, P ≤ 0.05; **, P ≤ 0.01; ***, P ≤ 0.001; ****, P ≤ 0.0001).

**Supplementary Fig. 10:**
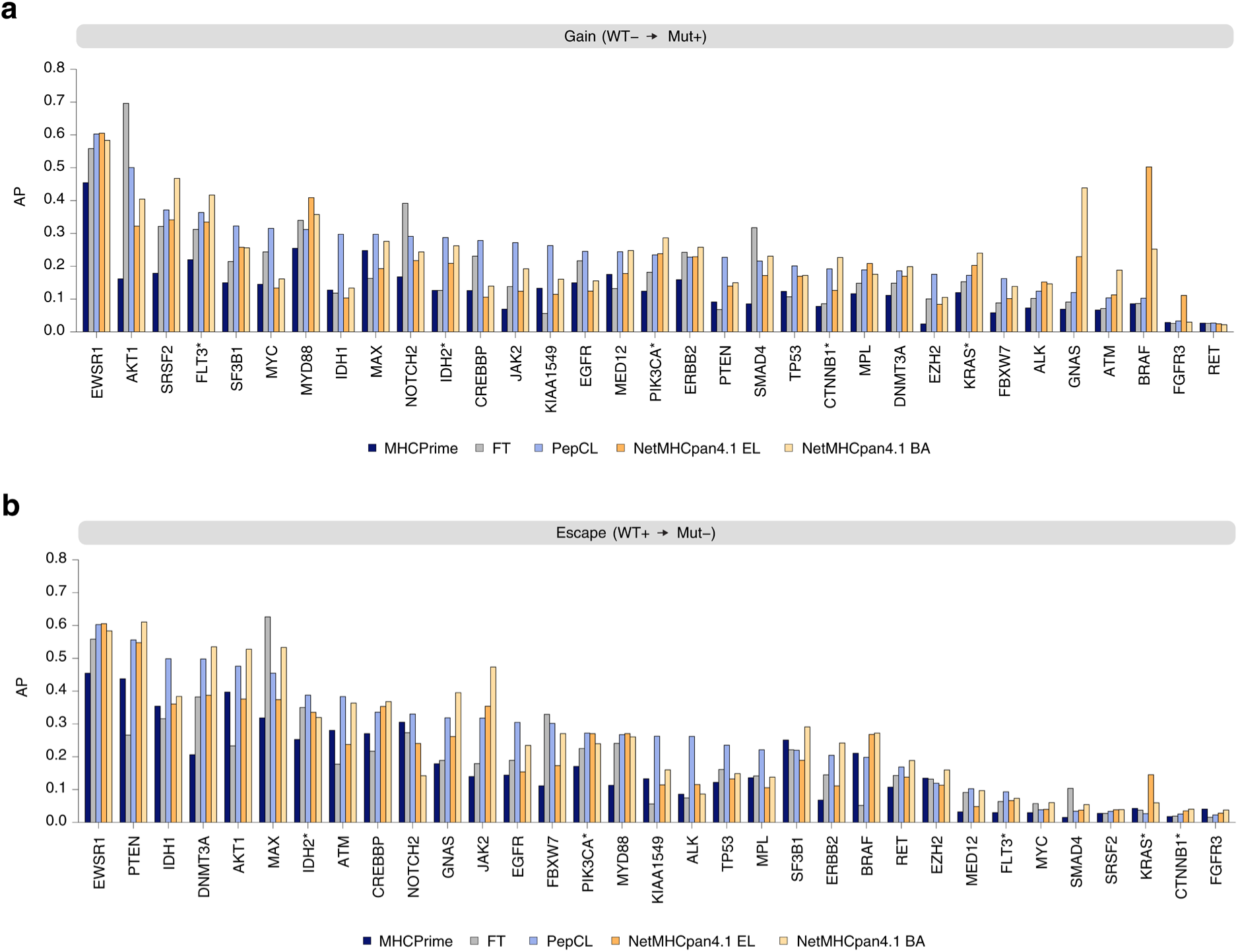
**Gene-held-out ESCAPE-seq mutation scanning**. **a)** AP for gain mutations in gene-held-out ESCAPE-seq experiments. **b)** AP for escape mutations in gene-held-out ESCAPE-seq experiments. For each held-out gene, model scores were computed for paired mutant and wild-type peptides, and differential mutation scores were calculated as mutant score minus wild-type score. AP was computed for the target mutation class against all negatives. The star (*) indicates genes that were held out together due to raw peptide overlaps (gene groups: KRAS (KRAS, KRAS-NRAS, HRAS, KRAS-KRAS-NRAS-NRAS-HRAS-HRAS, KRAS-NRAS-HRAS, NRAS-HRAS, KRAS-HRAS), FLT3 (FLT3, KIT, PDGFRA, FLT3-KIT-PDGFRA, FLT3-PDGFRA), IDH2 (IDH2, IDH2-IDH2), PIK3CA (PIK3CA, PIK3CA-PIK3CA), CTNNB1 (CTNNB1, CTNNB1-CTNNB1)).

## Notes

https://github.com/ntranoslab/MHCPrime

https://github.com/ntranoslab/PepCL

